# Iterative immunogen optimization to focus immune responses on a conserved, subdominant influenza hemagglutinin epitope

**DOI:** 10.1101/2025.11.04.686334

**Authors:** Daniel J. Marston, Akiko Watanabe, A. Brenda Kapingidza, Caitlin Harris, Elizabeth Van Itallie, Kapil Devkota, Jayani Christopher, Peiying Liu, Beena Pun, Brianna Rhodes, McKenzie Frazier, Kaitlyn Winters, Heman Burre, Pranav Kannan, Xiaoe Liang, Garnett Kelsoe, Kevin Wiehe, Rohit Singh, Daniel Wrapp, Masayuki Kuraoka, Mihai L. Azoitei

## Abstract

Designing effective vaccination strategies against genetically diverse viruses, such as HIV or influenza, is hindered by the ability of these pathogens to mutate and readily evade immune responses. While these viruses contain conserved regions that cannot easily mutate without affecting viral fitness, these sites are subdominant, meaning that they are not the primary target of antibodies elicited by natural infection or current vaccines. Here, we integrated recent advances in machine learning methods for protein structure prediction and design to engineer protein immunogens that focus immune responses on a conserved, subdominant HA epitope targeted by broadly cross-reactive influenza antibodies. Iterating between computation-guided optimization and *in vivo* analyses generated immunogens that faithfully displayed the target epitope and redirected humoral responses toward it upon vaccination. These results provide a blueprint for applying recent protein engineering approaches to immunogen design and may inform the development of broadly protective influenza vaccines.

## Introduction

Genetically diverse viruses, such as influenza, HIV, or coronaviruses, can rapidly mutate to evade immune responses^1^. The continuous evolution of these viruses has made the development of broadly protective vaccines against them challenging, as natural infection or existing vaccines typically provide protection only against the exposure strain or closely related variants^2–4^. Despite their enormous overall sequence diversity, the surface proteins of these viruses contain highly conserved regions that are usually critical for host receptor engagement and fusion with target cells^5,6^. Antibodies against these conserved epitopes have been isolated and demonstrate broad control over diverse viral variants, providing templates for “universal” vaccines that protect against multiple strains of a given virus^7–9^. However, conserved epitopes are typically subdominant, meaning that they are bound by only a small fraction of the elicited antibody responses following exposure to native viral proteins. Therefore, a significant goal of vaccine development across pathogens is to design immunogens that preferentially focus immune response on virus sites conserved in both current and emerging variants.

The trimeric hemagglutinin (HA) protein decorates the influenza virions and contains both the host receptor binding site (RBS) and the fusion machinery, which are essential for infection. The HA is the primary target of natural immunity against influenza and the main component of current seasonal influenza vaccines^10,11^. However, the majority of immune responses elicited by either infection or vaccination target antigenically variable sites on HA that can mutate without affecting viral fitness, eventually escaping immune responses^12^. This necessitates the annual update of influenza vaccines to match circulating strains, a process that can result in vaccines with variable efficacy and limited protection against emergent strains or those with pandemic potential, such as the recent avian H5N1 virus. To address these limitations, various strategies are currently being pursued to develop “universal” influenza vaccines that are broadly protective against diverse viral strains and do not require regular reformulation^13^. One such approach involves designing immunogens to preferentially elicit antibody responses that target conserved HA sites, such as the receptor binding site, the stem, or the more recently identified 220-loop at the interface of different HA head subunits^13^. Multiple human antibodies have been isolated against the 220-loop epitope, including FluA-20^7^ and S5V2-29^8^, which have been extensively characterized. These antibodies bind the majority of influenza A HAs and protect against diverse live virus challenges in animal models, including H3N2, H1N1, and H5N1^7,8,14,15^. While it would be beneficial to induce high titers of such antibodies by vaccination, neither current vaccines nor natural infections achieve this. This is likely due to the poor accessibility of the 220-loop on the native HA molecule and the presence of other immunodominant sites, like the HA head, that limit immune responses to the HA head interface. In the structures of pre-fusion HAs, the 220-loop is completely occluded and not accessible to antibodies. Still, it must become exposed at least temporarily through molecular breathing since both FluA-20 and S5V2-29 protect against challenges with live influenza viruses. Similar poorly accessible, subdominant epitopes that are targeted by broadly protective antibodies are also found on other viral proteins, like the HIV Envelope^16^, the coronavirus spike^17,18^, or Hepatitis C E1 E2 proteins^19^. Therefore, developing immunogens that elicit high titers of broadly cross-reactive antibodies against the 220-loop represents a stringent test case of our ability to rationally focus immune responses on desired epitopes and could have translational implications for developing a “universal” influenza vaccine.

Different vaccine design strategies have been employed to modulate the natural immunogenicity of virus proteins, aiming to enhance the accessibility of target epitopes and minimize undesirable off-target responses. The addition of glycosylation sites can be used to mask immunodominant regions and may consequently boost responses to typically subdominant sites^20^. An HA trimer immunogen that contained five additional glycosylation sites and aimed to boost immune responses to the RBS unexpectedly increased antibody responses to the HA head trimer interface. However, the elicited antibodies only bound H3 and H4 strains, so are likely not as broad as FluA-20 or S5V2-29^20^. Heterologous prime-boosting vaccination regimens that employ different immunogens can also focus antibody responses on shared epitopes^21–23^. However, these strategies are not always effective, can be difficult to generalize from one immunogen to another, and typically require multiple rounds of empirical testing. For example, the structure of a given immunogen limits the sites to which new glycosylation sites can be added^24^. At the same time, their occupancy or glycan type cannot be predicted, with shorter glycans providing limited coverage.

To improve the presentation of occluded viral epitopes, one effective strategy involves transplanting them in the conformation recognized by target antibodies onto unrelated “scaffold” proteins, resulting in a type of immunogen called epitope scaffolds ^25–27^. Beyond optimal structural epitope presentation, epitope scaffolds have the added benefit of a molecular environment devoid of distracting, immunodominant epitopes typically present on viral proteins. We and others have developed computational modeling and high-throughput screening approaches to graft structurally diverse epitopes onto non-viral proteins and showed that they elicit antibodies to the target epitopes through vaccination^27,28^. A major consideration of epitope scaffold design is that the grafted epitope accurately mimics the conformation recognized by target antibodies on native viral proteins to ensure that the antibodies they elicit can cross-react with the target virus. Besides the epitope conformation, it is important that the supporting scaffold does not inadvertently contain potent immunogenic regions that can reduce immune responses to the grafted epitope and instead focus them on undesired scaffold sites. Therefore, incorporating design approaches to rationally optimize epitope presentation and explicitly minimize antibody responses to the scaffold has the potential to advance the design of broadly protective vaccine candidates against conserved, but subdominant, viral epitopes.

In the last five years, machine learning algorithms have revolutionized protein structure prediction and design, enabling the rapid development of proteins with novel backbone structures and functions. These methods have the potential to enhance and accelerate the development of vaccine candidates. For epitope scaffold design, generative AI methods can produce scaffolds not found in nature that are customized to display complex epitopes^29–31^, while structure prediction algorithms can reveal with high confidence the conformation of the grafted epitope and inform ways to optimize it^32,33^. While still lagging in accuracy compared to protein prediction methods, the modeling of antibody-antigen complexes can also provide valuable information on epitope recognition by target antibodies^32,34^. Here, we integrated recently developed ML-based protein modeling methods into existing workflows to engineer epitope scaffolds that display the 220-loop of HA head, with the goal of eliciting antibodies like FluA-20 or S5V2-29 by vaccination (**Figure 1**). Given the unstructured loop conformation of this epitope, accurate prediction of its grafted structure was essential to guide iterative affinity improvements of engineered immunogens, resulting in high affinity binding to multiple target antibodies. Structural analysis revealed that the conformation of the epitope on the scaffold closely matches that recognized by target antibodies on the native HA. To increase the magnitude of elicited antibody responses targeting the grafted 220-loop upon vaccination, the scaffold framework was computationally redesigned to eliminate distracting immunogenic sites identified by modeling complexes between the epitope scaffold immunogen and representative antibodies elicited *in vivo*. These results provide a blueprint for applying novel ML-based protein prediction and design methods to enhance the rational design of immunogens that aim to focus antibody responses on conserved, subdominant viral epitopes, with the HA 220-loop as a test case. Our findings are also relevant for the development of universal influenza vaccines that aim to target the HA head trimer interface for broad protection.

**Figure 1.**
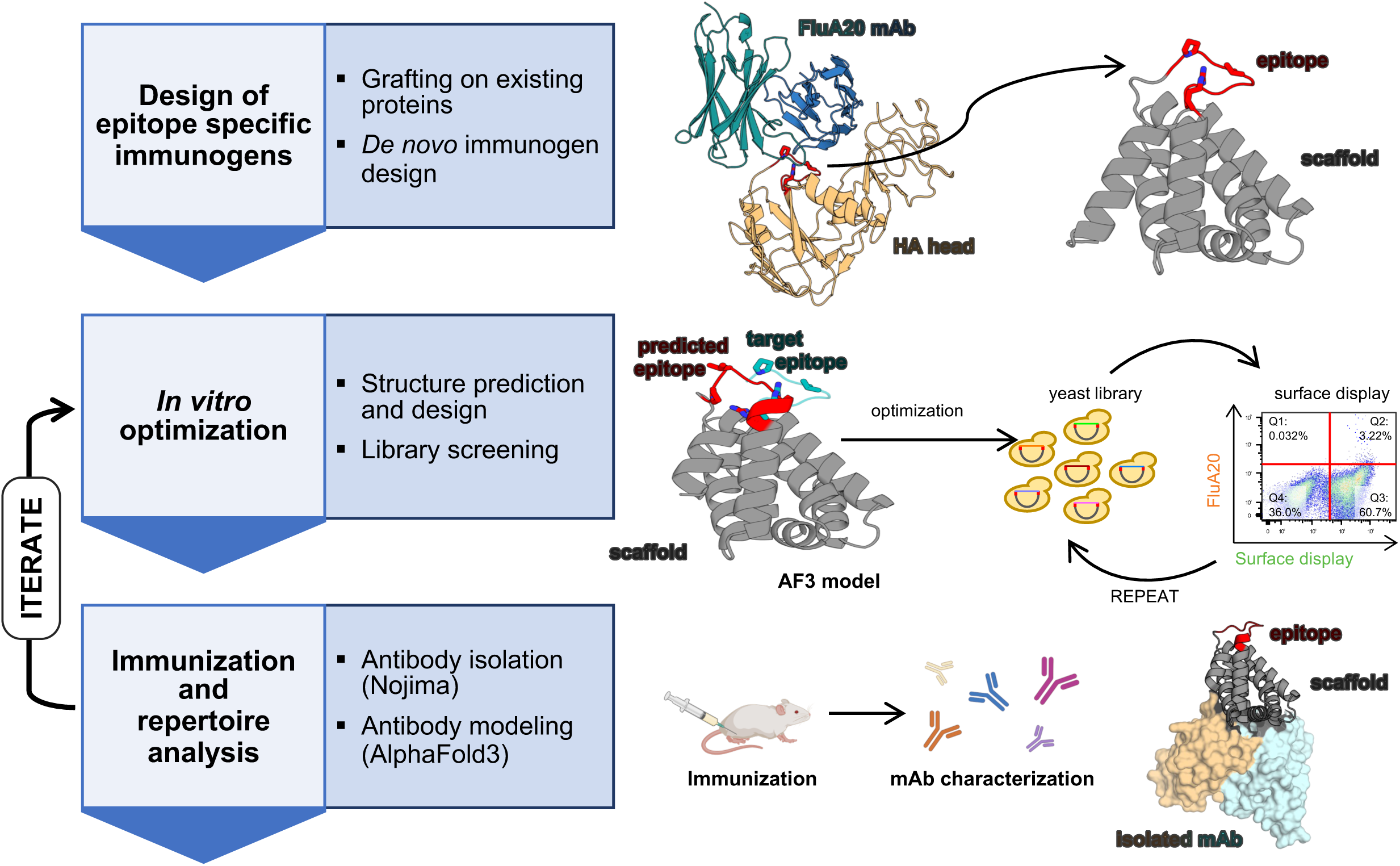
Immunogen design, characterization, and optimization pipeline. Workflow steps (*left*) are listed alongside associate techniques (*center* and *right*).

## Results

### Computational design of 220-loop epitope scaffold immunogens

The region spanning residues 220-229 of the influenza HA head, the 220-loop, forms part of the trimer interface (**Supplemental Figure 1**). The surface of the 220-loop that faces away from the receptor binding site (RBS) is highly conserved and targeted by multiple broadly cross-reactive antibodies, including FluA-20^7^, S5V2-29^8^, H5.31^14^, S1V2-58^8^, and D2 H1-1/H3-1^15^, with key contacts made with virus residues Pro221 and Val223 (**Figure 1**). These residues are occluded in the structure of the closed, trimeric prefusion HA head, likely limiting B Cell Receptor access and the elicitation of antibodies against this site in high titers. To improve the exposure and immune recognition of this epitope, we developed 220-loop epitope scaffolds immunogens. The 220-loop epitope is a long, flexible loop without well-defined structural elements, and therefore a challenging target to display on heterologous proteins. Because of this, the design process required multiple steps (**Figure 1**), including: (1) initial computational immunogen design; (2) multiple rounds of *in vitro* optimization; and (3,4) immunization followed by experimental and computational analysis of the induced antibodies. 220-loop epitope scaffolds were engineered using “backbone grafting” computational methods that we previously described^25,35^. In this approach, the target epitope replaces the backbone of an existing protein selected from a subset of ∼10,000 high-resolution proteins from the Protein Data Bank (**Figure 1**). This candidate scaffold library^35^ was queried computationally to identify proteins with exposed backbones where the N- and C-termini of the epitope could “land”, i.e. the ends of the epitope were compatible with the existing scaffold backbone. The epitope was then inserted into the scaffold backbone, replacing the native protein structure, and additional mutations were introduced to integrate the epitope into the scaffold and to ensure that the FluA-20 mAb could access the epitope in this new molecular context. This process identified one successful design, which we named HA220-1, built on the backbone of the VHS domain of the human Golgi-localized, γ-ear-containing, ADP-ribosylation-factor-binding protein 3 (GGA3)^36^. Transplantation of the 220-loop (HA residues 220-229) necessitated a significant redesign of the base scaffold, involving three mutations and ten insertions/deletions. The epitope replaced a region of the scaffold that was structurally different (RMSD=6.8Å between native and grafted epitope backbone). HA220-1 had weak binding to FluA-20 (**Supplemental Figure 2**), but the binding was mediated by contacts located in the epitope, since mutations known to knock down FluA-20 binding on HA head had the same effect on HA220-1.

### In vitro optimization of 220-loop epitope scaffolds for increased affinity to target antibodies

The limited binding of FluA-20 mAb to HA220-1 suggested that its epitope presentation is suboptimal. To guide affinity improvements and gain insight into the epitope conformation, the HA220-1 structure was modeled using AlphaFold2 (**Figure 2A**), which had been recently released at the time. The resulting models suggested that the grafted epitope adopted a conformation distinct from that recognized by FluA-20 in the structure of its complex with the native HA head (RMSD = 4.5Å). To improve the epitope presentation and increase antibody affinity, we employed *in vitro* affinity maturation using yeast surface display and sampled all the possible amino acid combinations at groups of five residues that connect the grafted epitope to the scaffold (**Figure 2B**). After three selection rounds by FACS using decreasing concentrations of FluA-20 (**Figure 2B**), we identified a new epitope scaffold variant, HA220-2, with improved binding to FluA-20 (*K_D_*=978 nM, **Supplemental Figure 2B**), and that also recognized 220-loop antibodies S1V2-58 and D2 H1-1/H3-1, albeit with micromolar affinities (8.8 μM and 2.7 μM, respectively, **Supplemental Figure 2B**). In previous work^35^, we showed that binding to diverse antibodies targeting the same epitope *in vitro* is a key marker for correct epitope presentation and is essential to ensure that the correct antibodies are elicited *in vivo* by vaccination.

**Figure 2.**
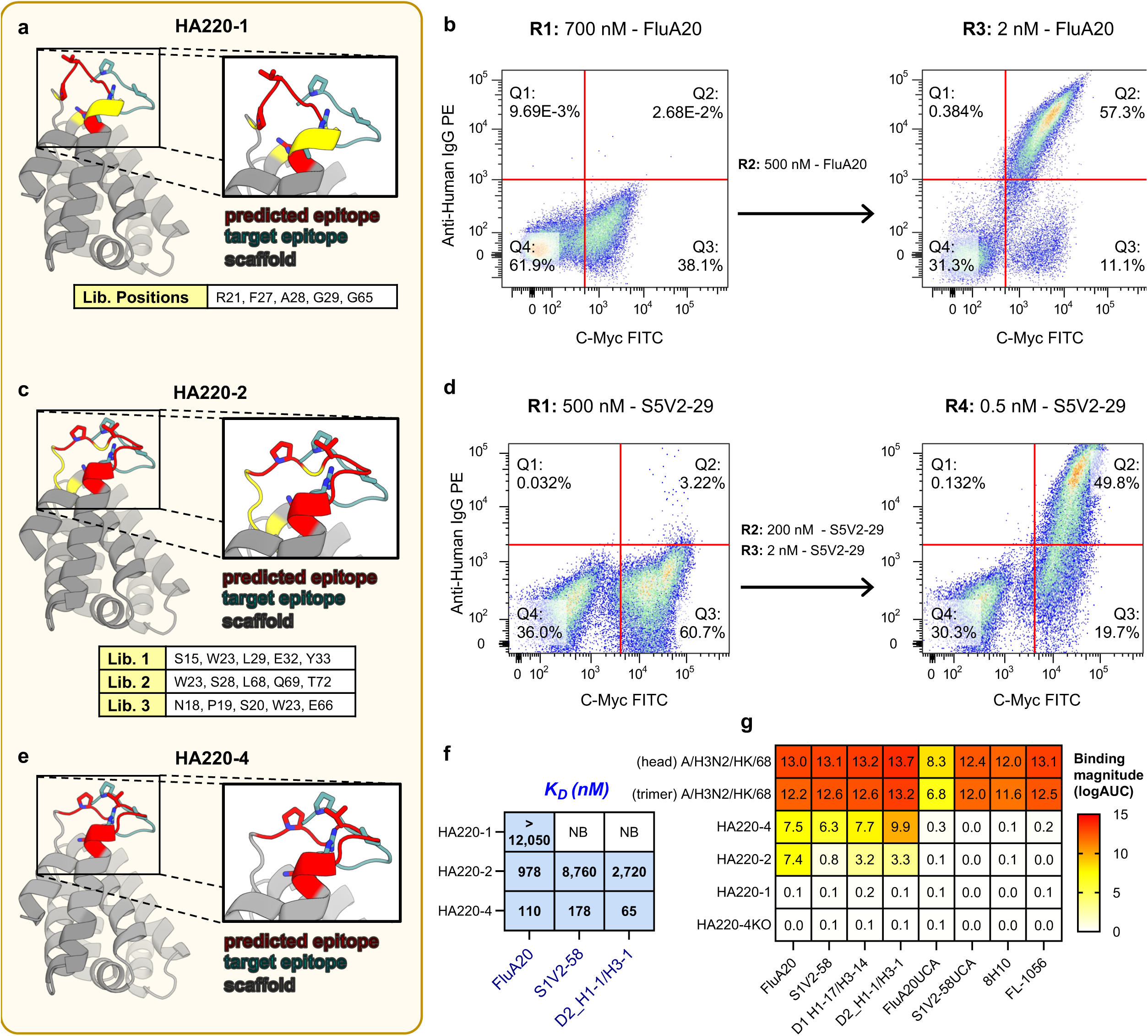
Development and characterization of first generation 220-loop epitope scaffolds. **(a)** AlphaFold2 model of HA220-1 (scaffold backbone in *grey*; predicted epitope conformation in *red*; target epitope conformation in *blue*). Epitope residues equivalent to P221, V223 and R229 (H3 numbering) shown in sticks. Sites diversified for *in vitro* optimization are marked in yellow and listed in the table. **(b)** FACS screening of HA220-1 libraries. R1-R3 describe the selection round number, the antibody used, and the labeling concentration. **(c)** AlphaFold2 model of HA220-2 isolated from *(b)* colored as in *(a)*. **(d)** FACS screening of HA220-2 libraries labeled as in *(b)*. **(e)** AlphaFold2 model of the HA220-4 immunogen isolated from *(d)* colored as in *(a)*. **(f)** Binding affinities of HA220 variants to 220-loop mAbs determined by BLI. NB= not binding **(g)** ELISA binding data of HA220 variants, A/Hong Kong/1/1968 (HK/68) trimer and head domain to target mAbs and their unmutated common ancestors (UCAs). HA220-4KO contains epitope mutations known to abrogate binding.

Therefore, to further improve the recognition breadth of HA220-2, we screened three new libraries with an alternative 220-loop binding antibody, S5V2-29, sampling additional combinations of sites adjacent to the epitope (**Figure 2C, D**). This led to the identification of a new variant, HA220-4 (**Figure 2E**), which shows further improved affinity for FluA-20 and also has nanomolar affinity for S1V2-58 and D2 H1-1/H3-1 mAbs, both of which bind to the 220-loop but use different VH genes, demonstrating significantly enhanced breadth (**Figure 2F, G, Supplemental Figure 2B**). Importantly, HA220-4 did not bind to 8H10 or FL-1066, two antibodies that engage the 220-loop but have limited breadth compared to FluA-20 or S5V2-29^20^. AlphaFold modeling showed that, compared to the HA220-1 model, the epitope conformation on H220-4 was expected to be closer to that of the 220-loop on the HA head (RMSD=4.1 Å), linking improved antibody recognition with a better predicted epitope mimicry.

### In vivo characterization of first generation 220-loop immunogens

Next, we assessed the immunogenicity of HA220-2 and HA220-4 by measuring their ability to preferentially elicit 220-loop antibodies in animals with pre-existing immunity to influenza. Mice were first primed with the HA head domain from A/H3N2/Hong Kong/1968 (HK/68 trimer) adjuvanted in alum, to simulate pre-existing immunity, and then boosted twice, 28 and 56 days later, with either HK/68 HA head or soluble forms of HA220-2 and -4 (**Figure 3A**). As expected, all animal sera reacted with HK/68 head, with the highest reactivity in the animals immunized with HK/68 three times (**Figure 3B**). While these sera showed some reactivity with other H3N2-derived HA heads, they did not bind any HA heads from other influenza A subtypes like H1, H5, or H7 (**Figure 3B**). Interestingly, some of the responses elicited by HA head immunization competed with the binding of S5V2-29 to the HK/68 head (**Figure 3C)**, suggesting the elicitation of antibodies that target an overlapping epitope with that of S5V2-29. However, given the lack of observed breadth, these antibodies are likely functionally distinct from S5V2-29 and may be more similar to 8H10 or FL-1066^20^. This suggests that natural HA heads do not induce detectable levels of broad antibodies against the 220-loop, and that their molecular environment contains other immunodominant regions around this epitope that further limit the elicitation of FluA-20 antibodies. Sera from animals boosted with the epitope scaffolds also showed limited breadth, except in a few animals (5 out of 24) that had reactivity to HA head domains from H5N1 strains (**Figure 3B**). Responses to the grafted epitope, as measured by the difference in binding to the epitope scaffold immunogen and to the original parent scaffold that did not contain the epitope, were limited in groups boosted with HA220-2 or HA220-4, accounting for 25% to 35% of total titers (**Figure 3D**). Taken together, these results showed that HA220 immunogens have a limited ability to focus responses on the 220-loop epitope and that they contain other, more immunogenic sites.

**Figure 3.**
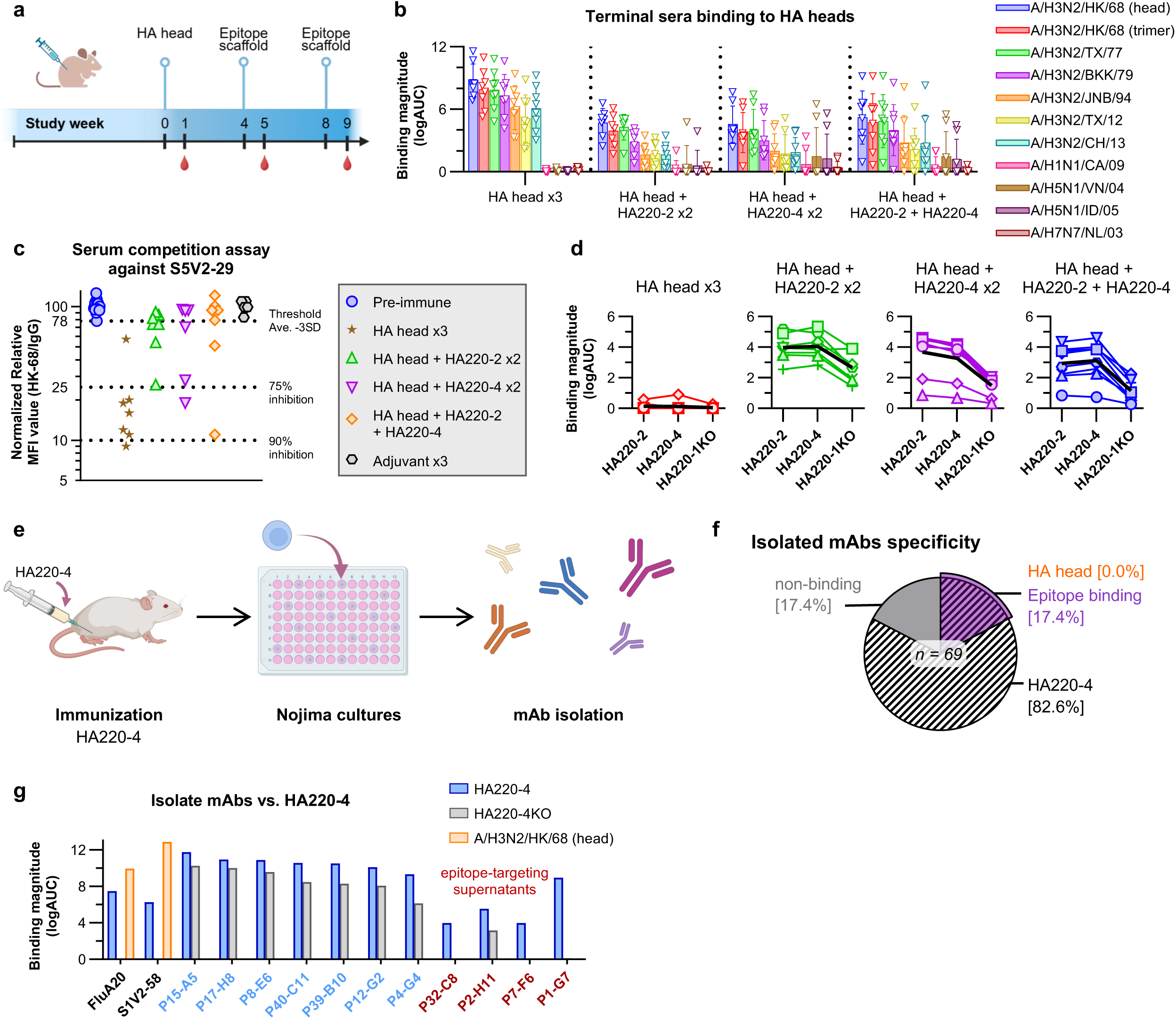
Immunogenicity of HA220-4. **(a)** Immunization protocol with HA220-2 and -4 in the context of preexisting immunity induced by the HA head domain of A/Hong Kong/1/1968 (H3N2). **(b)** Sera binding to HA heads from the indicated influenza strains. X-axis lists the four different immunization regimens. Y-axis values are log(AUC) of ELISA titration binding curves. **(c)** Sera blocking of S5V2-29 mAb binding to HK/68 HA head. The y-axis indicates the normalized MFI binding relative to the maximal binding of S5V2-29 mAb to HK/68 HA head in the absence of competitor. Dotted lines mark 75 and 90% inhibition of binding. Each data point represents an individual animal from the respective immunization group. **(d)** Sera binding to the epitope scaffolds used in the immunization and a KO version. **(e)** Schematic of Nojima culture analysis from mice vaccinated once with HA220-4. **(f)** Binding specificity of 69 mAbs isolated and characterized by Nojima. **(g)** ELISA binding (log(AUC)) of recombinant isolated mAbs to HA220-4 and HA220-4KO. Binding of FluA-20 and S5V2-29 mAbs to HK/68 head were included as control. **(h)** AlphaFold3 models of representative mAbs from *(g)* in complex with HA220-4 (scaffold backbone: *grey*; epitope: *red*; mAb *cyan*).

To gain a deeper understanding of the immune response triggered by the HA220-4 immunogen, which did not contain significant antibody titers to the 220-loop, we conducted Nojima culture analysis^37^ to characterize the elicited antibody repertoire. Individual B cells were isolated from germinal centers within the inguinal lymph nodes of mice that received one shot of the HA220-4 immunogen (**Figure 3E**). Individual cells were cultured in Nojima cultures, class switched to secrete IgG antibodies, and the resulting supernatant was assessed for binding to a panel of diverse HA heads. After culture, we screened 69 clonal IgG Abs in culture supernatants, of which 57 (82.6%) bound the HA220-4 immunogen. Differential binding to HA220-4 and to an epitope knockout version that eliminated FluA-20 binding revealed that only 6 out of the 57 HA220-4 binding supernatants engaged the epitope, with the majority of the elicited antibodies likely targeting other scaffold regions (**Figure 3F**). However, none of the epitope targeting antibodies elicited by the immunogen cross-reacted with natural HA heads (**Figure 3G**), suggesting that they may require interactions with the scaffold that are not present on HA or that they recognize an epitope conformation distinct from the native one.

To identify regions of the HA220-4 that are targeted by elicited antibodies and may distract from the epitope, we first selected a panel of 11 Nojima antibodies, expressed them recombinantly, and confirmed their binding by ELISA (**Figure 3C**). We modeled the interaction of these antibodies with HA220-4 using AlphaFold3 (iPTM>0.6) and identified several potential scaffold regions that may be immunogenic (**Supplemental Figure 3**).

### Epitope scaffold optimization by deep scanning mutagenesis

To address the limitations of the 220-loop epitope scaffolds, we next pursued parallel strategies to: 1) further improve the epitope presentation of HA220-4 to achieve higher affinity for target antibodies; and 2) modify the surface of HA220-4 to reduce off-target scaffold responses. Since we previously optimized HA220-4 through directed libraries that sampled combinations at 16 sites around the epitope, we next took an alternative approach by examining the effect of single substitutions at 63 positions located as far as 15 Å from the epitope (**Figure 4A**). This library aimed to capture beneficial mutations away from the epitope that could improve the epitope structure through allosteric effects or improve the overall scaffold stability. This library was sorted by FACS in parallel for binding to three different 220-loop binding antibodies, FluA-20, D2 H1-1/H3-1, and S1V2-58 to capture mutations beneficial for engaging diverse antibodies (**Figure 4B**). The DNA of selected binding clones was sequenced using Next Generation Sequencing, and the library frequency of each mutation was determined before and after antibody selection to assess whether it was beneficial or detrimental to binding (**Figure 4C**). The mutations at each site that show neutral or positive enrichment across all three antibodies were calculated to identify immunogen positions that restrict antibody binding (**Figure 4D**). This library included previously optimized sites (Arg21, Thr27, Thr28, Leu29, and Val65 from the initial screen and Trp23, Thr28, Met68, Asn69 and Val72 from the secondary screen). The single-site mutagenesis data showed that these sites did not tolerate multiple changes, particularly those within the epitope (residues 21-30) and at amino acid Asn69, which lies directly underneath the epitope, suggesting that they had been previously successfully optimized (**Figure 4D**). Interestingly, FluA-20 was more sensitive to changes than the other two antibodies. Initial inspection of the data did not reveal any obvious substitutions that enhanced binding to all three antibodies. Furthermore, three changes, which showed the largest positive effects in FluA-20 (HA220-4-SV1, 2, and 3), failed to improve binding (**Figure 4E**).

**Figure 4.**
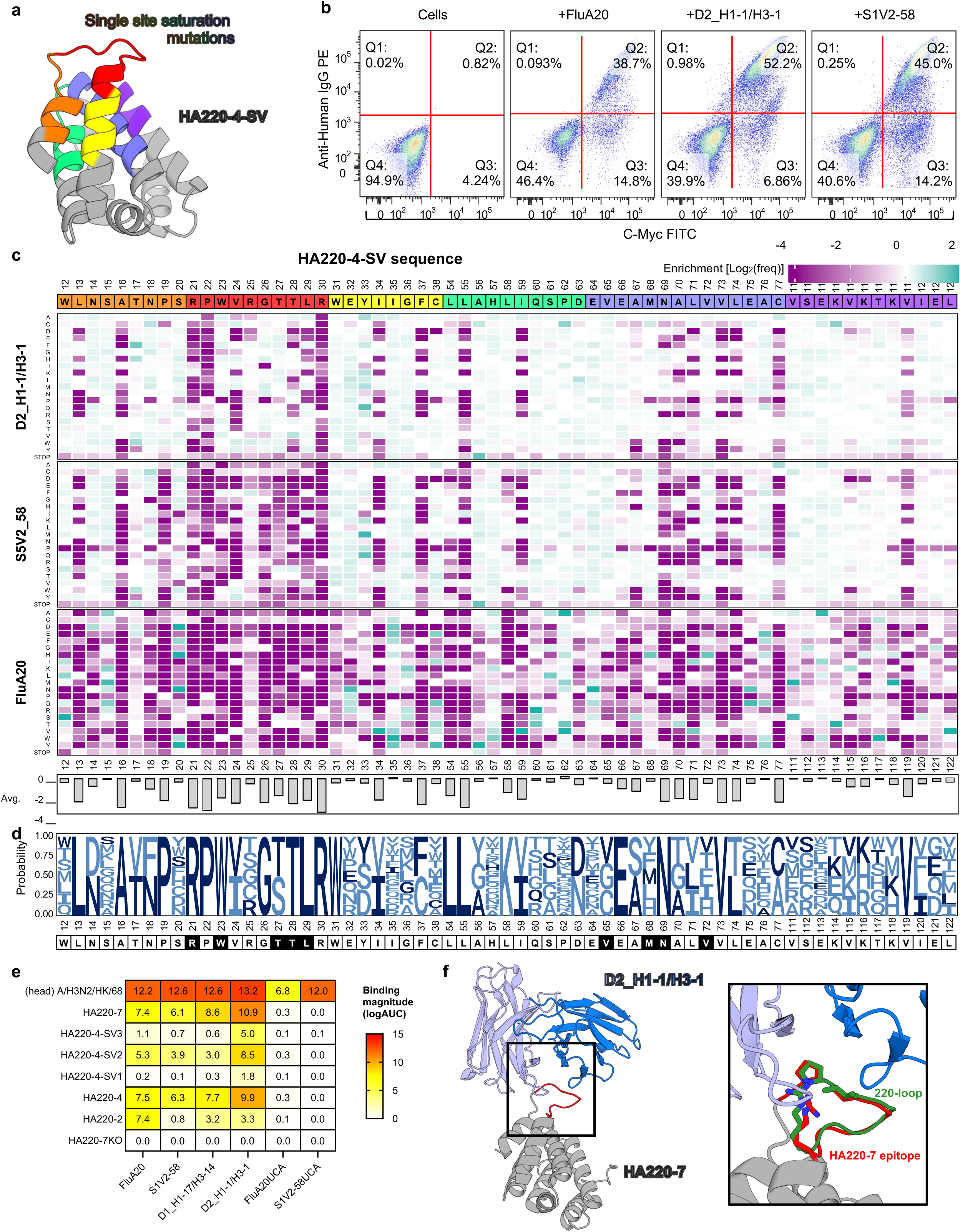
HA220-4 optimization by single-site saturation libraries. **(a)** AlphaFold2 model of HA220-4 (epitope scaffold: *grey*; predicted epitope conformation: *red*). Mutated sites are colored, with their sequences provided in *(c)*. **(b)** Library analysis by FACS to select clones binding to FluA-20 *(left)*, D2 H1-1/H3-1 *(middle)*, or S1V2-58 *(right)* mAbs, respectively. **(c)** Effect of every mutation on binding to the three individual mAbs calculated as enrichment levels (*purple*: unfavorable; *blue*: favorable; *white*: neutral). The HA220-4 reference sequence is shown at the top and colored as in *(a)*. Average enrichment for mutations at one position across all three mAbs is shown in the lower panel. **(d)** Logo plot of substitutions that have a neutral or positive effect on binding to all three mAbs. **(e)** Binding by ELISA of HA220 immunogens and their optimized variants (SV1-3) to target antibodies and their unmutated common ancestors (UCAs). HA220-7KO contains epitope mutations known to abrogate mAb binding. HA head domains from A/Hong Kong/1/1968 (H3N2) are included as reference. **(f)** Crystal structure of the D2 H1-1/H3-1 mAb (*blue*) in complex with HA220-7 (*grey*, epitope in *red*). Inset shows details of the epitope/mAb interaction with residues equivalent to P221, V223 and R229 (H3 numbering) shown in sticks. The epitope conformation induced by D2 H1-1/H3-1 on the HA head domain from the A/H3N2/TX/12 strain (PDBID:6XPR^15^) is shown in *green*.

To identify mutations that enhance the binding of HA220-4 to the three target antibodies, we developed a computational pipeline to efficiently predict the effect of 2-5 mutations on antibody recognition, based on the high-throughput screening data. We used Protein Language Models (PLMs) to identify beneficial combinations of mutations. PLMs are computational models trained on millions of protein sequences to learn the statistical patterns underlying protein structure and function. These models convert protein sequences into high-dimensional numerical representations called embeddings, which encode structural and functional constraints learned from natural protein evolution. We trained three different PLMs (ESM-2, ESM-1v, and Raygun) on our single-point mutation data, enabling them to predict antibody binding affinities from sequence information alone. By selecting only mutations ranked highly by all three models, we identified candidates that balanced improved antibody binding with maintenance of proper protein folding. This protocol led to the development of 20 HA220-4 versions, each containing 2-5 mutations, which were expressed recombinantly and assessed for their ability to bind antibodies. Sixteen variants expressed as soluble proteins, with nine showing measurable binding to FluA-20 (**Supplemental Figure 4A**), a relatively low success rate considering that all variants contained mutations predicted to be favorable for binding individually. Nevertheless, three novel variants had higher affinity than HA220-4, with one, called HA220-7, that included two mutations away from the epitope (I35V; P62A), showing markedly higher expression and ∼10-fold higher avidity for FluA-20 by ELISA (**Figure 4E**). These improvements facilitated the determination of a high resolution (2.4Å) crystal structure of HA220-7 bound by the 220-loop targeting antibody D2 H1-1/H3-1 (**Figure 4F, Supplementary Table 1**). This structure revealed that the bound epitope on HA220-7 adopted the same conformation as that recognized by the antibody in the context of the HA head (backbone RMSD= 0.8Å), illustrating the successful transplantation of the epitope loop on this scaffold (**Figure 4F**).

### Redesign of immunodominant sites to focus responses on epitope

To reduce the immunogenicity of the HA220 base scaffold shared by all the engineered immunogens, its surface was redesigned to eliminate binding of off-target antibodies (**Figure 3G**). Surface residues were altered using ProteinMPNN^31^, an ML-based protein design tool that takes a backbone structure as input and generates a protein sequence predicted to adopt the same fold. We used a model generated by backbone grafting as described above as the input for ProteinMPNN, given that it displayed the 220-loop structure bound by FluA-20, and produced hundreds of different sequences, which were screened using AlphaFold to ensure correct epitope conformation. Fifteen total designs were tested experimentally. Five of them maintained the key epitope contact residues (Pro221, Val223, Arg230) fixed, while the other 10 designs kept unchanged both the epitope residues as well as scaffold residues previously identified by screening to be favorable for antibody binding (residues 21-30 and Asn69); in both cases, the rest of the protein surface was allowed to change during ProteinMPNN design. The tested designs had 58-93% sequence identity with each other and 43-57% identity with the natural scaffold (**Figure 5A**). Although Alphafold2 predicted correctly folded loops for all the designs tested, only 1 of the designs, H220-M4, bound to FluA-20, although several bound weakly to other 220-loop antibodies (**Figure 5B, Supplemental Figure 4B)**. H220-M4 was from the design group that preserved more residues and bound all the 220-loop antibodies tested, albeit weakly (**Figure 5B**). Importantly, H220-M4 did not interact with any of the isolated antibodies elicited by H220-4, suggesting that immunodominant sites present on H220-4 and the native scaffold have been modified (**Figure 5C**). Therefore, this approach has successfully altered the immunogenicity of the HA220-M4 scaffold, compared to the scaffolds employed in HA220-4 and HA220-7.

**Figure 5.**
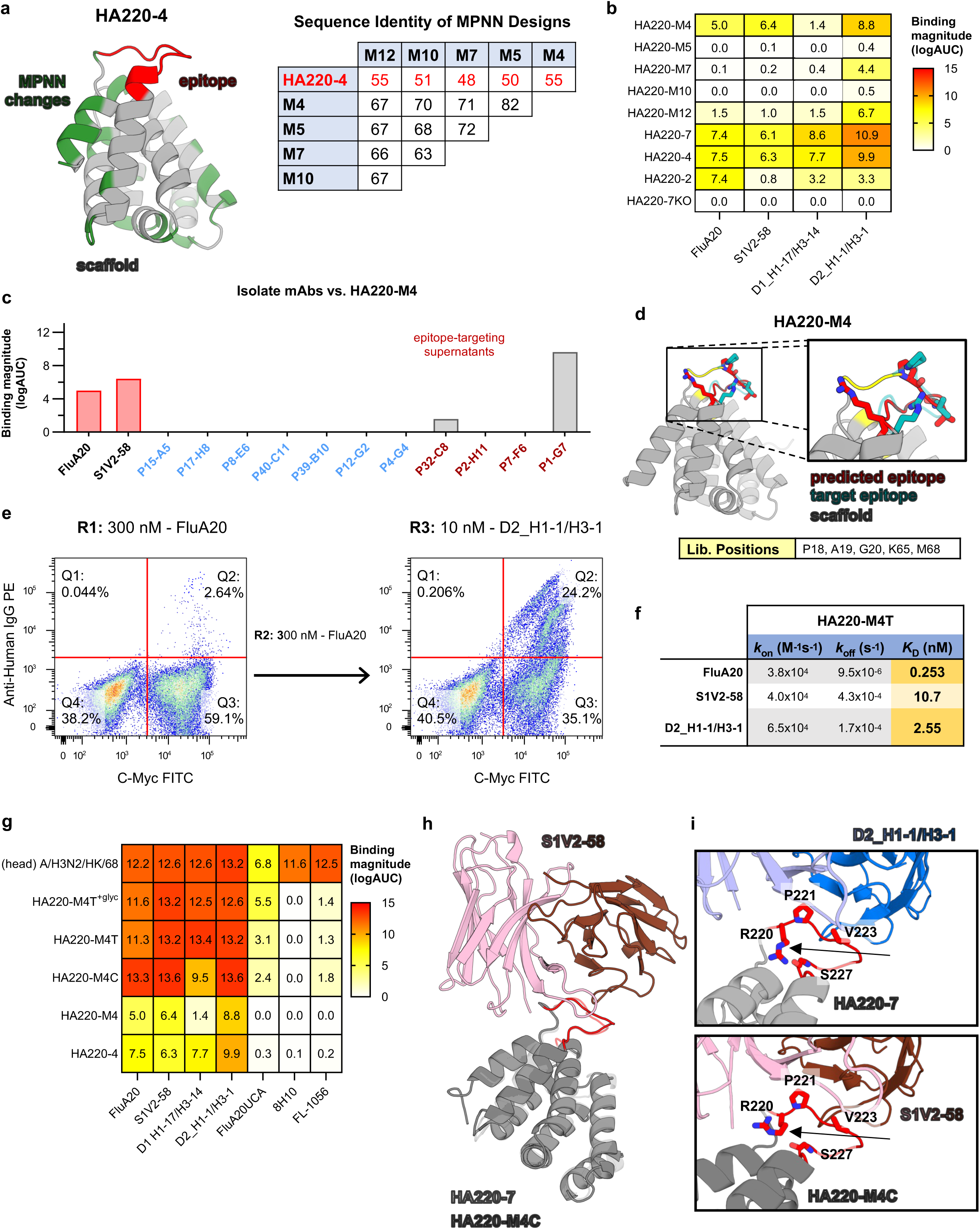
Iterative optimization of HA220 immunogens by computational modeling and high-throughput library screen. **(a)** *Left*: AlphaFold2 model of HA220-4 (scaffold: *grey*; epitope: *red*) with surface sites redesigned by ProteinMPNN in *green*. *Right*: Sequence identity of selected ProteinMPNN designs compared to HA220-4. **(b)** ELISA binding of ProteinMPNN design HA220-M4 to target antibodies, with previous HA220 versions shown for comparison. **(c)** ELISA binding of HA220-M4 to scaffold specific mAbs isolated by HA220-4 immunization. **(d)** AlphaFold2 model of HA220-M4 (scaffold: *grey*; epitope: *red*). Reference epitope conformation bound by FluA-20 on the HK/68 HA head is shown in *blue*. Sites diversified for *in vitro* optimization are marked in yellow and listed in the table. **(e)** FACS screening of HA220-M4 libraries. Selection rounds (R1-3), selection mAb and working concentration are indicated. **(f)** Binding affinities of the HA220-M4T and C variants isolated from *(e)* to selected 220-loop mAbs. **(g)** ELISA binding of HA220-M4 variants to target antibodies and their unmutated common ancestors (UCAs). The A/Hong Kong/1/1968 HA head domain and HA220-4 are included for reference. **(h)** Crystal structure of the S1V2-58 mAb (*blue shades*) bound to HA220-M4C (*grey,* epitope in *red*), overlayed with the structure of HA220-7 from Figure 4f (*light grey*). **(i)** Inset shows details of the epitope/mAb interactions (D2 H1-1/H3-1 + HA220-7, *upper*; S1V2-58 + HA220-M4C, *lower)*, with the R220 residues indicated by black arrows. Key epitope residues are numbered (H3 numbering) and shown in sticks.

Given that HA220-M4 had a novel sequence and it only bound 220-loop mAbs weakly, we undertook one more *in vitro* evolution round to further improve its affinity for diverse 220-loop antibodies. We varied residues adjacent to the N-terminus of the epitope that had not previously undergone selection and additional residues that supported the epitope (18, 19, 20, 65 and 68). Libraries were sorted twice with FluA-20 and then with D2 H1-1/H3-1 to isolate binders with breadth to multiple 220-loop antibodies (**Figure 5D, E**). The top immunogens isolated from this library, called HA220-M4C and HA220-M4T, had the highest binding affinities of any HA220 immunogen to multiple 220-loop monoclonals (HA220-M4T/FluA-20*, K_D_*=0.25nM, **Figure 5F, Supplemental Figure 2B**). They were also the only designs to bind the inferred Unmutated Common Ancestor (UCA) of FluA-20, suggesting they may more easily engage naïve B cells that can evolve and secrete these types of antibodies *in vivo* (**Figure 5G**). A 3.9 Å crystal structure of HA220-M4C in complex with the 220-loop antibody, S1V2-58 (**Figure 5H, Supplementary Table 1**), showed that its epitope adopted a very similar shape to that of native HA head (RMSD=0.5 Å) and HA220-7 (RMSD=0.5 Å) (**Figure 5H**). One major structural difference that may explain the weaker binding of HA220-7 relative to HA220-M4C, is that the arginine at the equivalent position to 220 in the HA 220-loop formed a hydrogen bond with the backbone carbonyl of position 227 in H220-7, while it pointed outside, more similarly to the conformation observed on HA head, in HA220-M4C (**Figure 5I**). This intra-epitope loop interaction in HA220-7 may render the epitope more rigid, limiting the ability of 220-loop antibodies to induce the optimal binding conformation.

### In vivo characterization of second generation 220-loop immunogens

Given their improved binding characteristics, we then characterized the immunogenicity of HA220-M4T and HA220-M4C *in vivo*. To further reduce potential off-target scaffold responses, glycosylation sites were introduced into the scaffolds, 2 into HA220-M4C and 4 into HA220-M4T, resulting in variants called HA220-M4C^+glyc^ and HA220-M4T^+glyc^ (**Figure 6A**). In a Nojima culture experiment, like the one initially conducted in our HA220-4 immunization experiment (**Figures 3E, 6B**), HA220-M4T^+glyc^ showed markedly improved immunogenicity by eliciting diverse antibodies against the 220-loop epitope that cross-reacted with native HA. After one immunization with HA220-M4T^+glyc^, of the 2107 isolated antibodies from five mice, 883 bound the HA220-M4T^+glyc^ immunogen. Of these, 6.9% targeted the epitope, as evidenced by binding to HA220-4 scaffold that contains the 220-loop in a different scaffold context (**Figure 6C**). Importantly, 62% of these epitope specific antibodies, or 4.3% of the total isolated HA220-M4T^+glyc^ reactive antibodies, also bound the HA head domain, with some engaging the full HA trimer, indicating that in four out of the five mice, we elicited bona fide 220-loop antibodies that can access their target on native influenza proteins. To further characterize these antibodies, the VDJ regions of the Nojima culture B cells from 3 mice were sequenced. Within these sequences, we identified multiple members of four B cell clones with distinct immunogenetics that bound to the 220 loop on native HA (**Supplemental Figure 5**). Selected antibodies from these clones were purified and showed binding to HK68 HA heads and HA trimers **(Figure 6D).** Interestingly, HA220-M4T bound to the inferred UCAs of some of these clones, while HA220-M4C did not, which may explain why only HA220-M4T was able to induce these types of antibodies. In the largest clone (6_18) all lineage members contained a single residue change, L46V in the light chain. This single mutation was sufficient to provide HA head binding when introduced into the 6_18_UCA (6_18_UCA(LV), **Figure 6D).**

**Figure 6.**
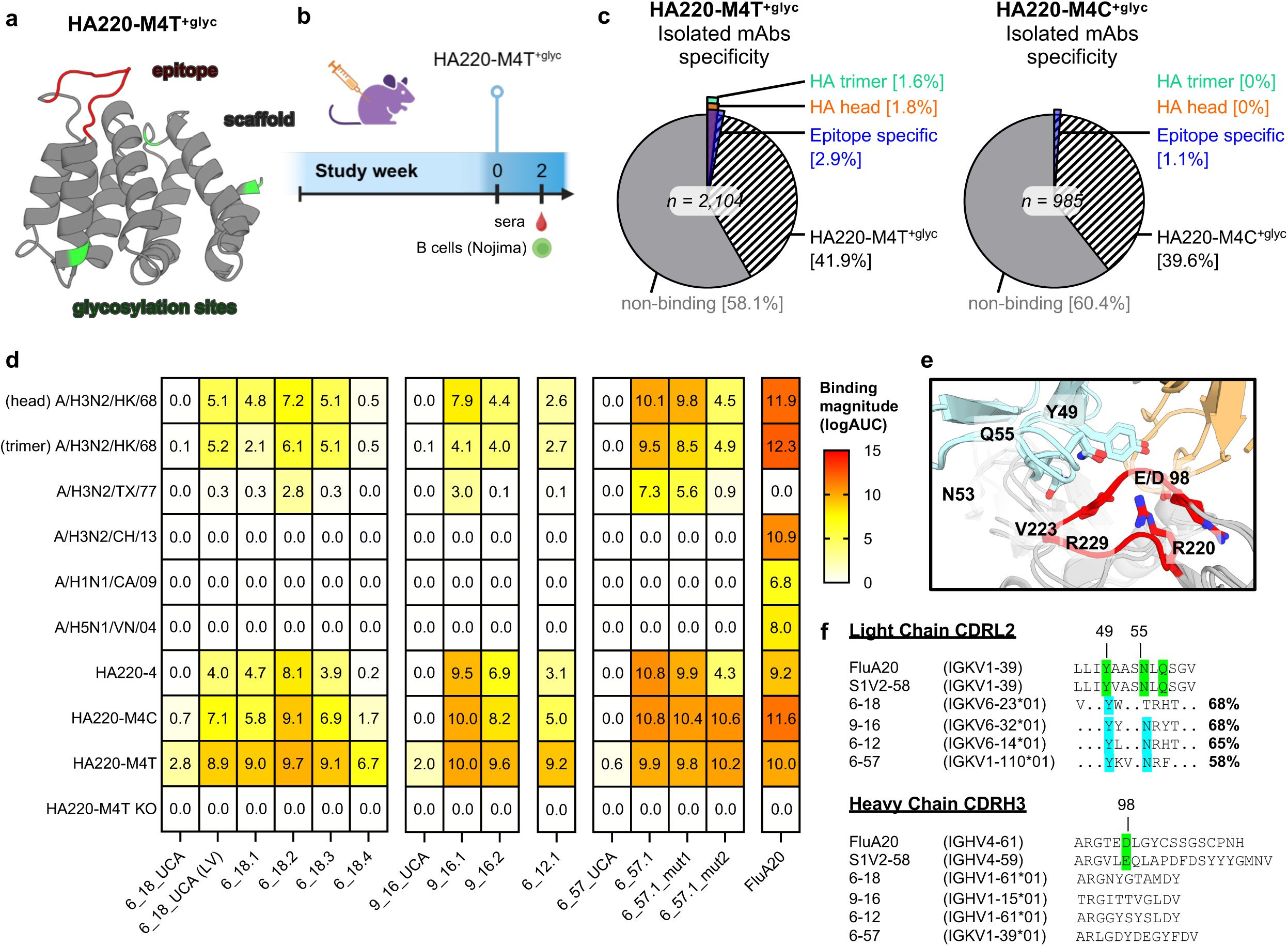
Optimized HA220-M4T^+glyc^ immunogens elicit epitope-specific antibodies. **(a)**AlphaFold2 model of HA220-M4T^+glyc^ epitope scaffold (*grey*), with glycosylation sites labelled (*green*). **(b)** Schematic of Nojima cultures analysis from animals vaccinated once with HA220-M4T^+glyc^. **(c)** Binding specificity of antibodies isolated and characterized by Nojima cultures after a single immunization with HA220-M4T^+glyc^. **(d)** ELISA binding (log(AUC)) of recombinant isolated mAbs to indicated HA heads and scaffolds. Binding of FluA-20 mAb included as control. **(e)** Overlay of crystal structures of mab D1 H1-17/H3-14 in complex with A/Texas/50/2012 (PDBID 6XPQ) and mab FluA-20 in complex with A/Hong Kong/1/1968 (PDBID 6OCB) with key interacting residues shown in sticks. Mab light chains in *cyan,* heavy chains in *orange,* and the 220 loop in *red.* **(f)** Alignment of CDRL2 (*upper)* and CDRH3 (*lower*) loops from FluA-20, S1V2-58, and germline sequences of isolated clones.

Although antibodies elicited by HA220-M4T^+glyc^ bound the HA of A/H3N2/Hong Kong/1968, which contains the same 220-loop sequence that was grafted, no binding was observed to heterologous HAs, like those derived from H1N1 or H5N1 strains. To uncover antibody features that may be important for breadth, we compared the immunogenetics of the elicited mouse antibodies with canonical 220-loop human ones. The broadest 220-loop human antibodies are typically encoded by VK1-39 in the light chain which is often paired with VH4-59 or VH4-61. These antibodies contact the 220-loop through conserved residues Tyr49, Asn53 and Gln55 in the CDRL2 loop and an acidic residue at position 98 within a long (20+ residues) CDRH3 loop **(Figure 6E).** No close relative of human VK1-39 exists in the mouse genome, the closest is IGKV10-94 at 75.8% identity. However, the antibody clones we isolated had Tyr49 and Asn53 as germline encoded residues (**Figure 6F),** and residue Leu46, which was key for clone 6_18 binding to HA head, is directly adjacent to these contacts suggesting these conserved residues may be important for the binding interaction. Additionally, the D and J genes encoded in the mouse genome dramatically reduce the likelihood of longer CDRH3 loops and this makes it less likely that an acidic residue at position 98 of the CDRH3 loop would orient towards the 220 loop correctly. To test whether either VK1-39 or the CDRH3 differences were key limiting features of this mouse model, we took our strongest trimer binding antibody (6-57.1) and modified it to incorporate these key features. The first modification was to mutate the CDRL2 loop so that it contained the key conserved residues, Gln55 and a small residue at position 50 (mut1). The second was to mutate the CDRH3 loop at position 98 to an acidic residue and combine this with the CDRL2 changes (mut2). Mutating the CDRL2 or position 98 had little effect on the breadth of binding, suggesting that these were not sufficient to recapitulate the broadest human 220 loop binding antibodies. It may be that the short mouse CDRH3 prevents these antibodies from recapitulating the full binding breadth of the human 220-loop-binding antibodies. However, although we were unable to achieve the fullest breadth, these results show that the iterative sequence optimization of HA220 immunogens to improve epitope conformation and minimize undesired scaffold responses led to the elicitation of antibodies that recognize the 220-loop on native HA molecules and share some features with canonical human antibodies against this site.

## Discussion

Focusing immune response onto conserved, typically subdominant, viral epitopes is a major focus of vaccine development efforts targeting antigenically variable pathogens, such as HIV, influenza, or coronaviruses. In many cases, such epitopes are occluded and poorly accessible to BCRs on the target viral proteins and surrounded by other immunodominant variable regions that attract the majority of elicited immune responses. For example, the stem helix and S^816–834^ (formally fusion peptide) epitopes on coronavirus spikes and the MPER region of the HIV Env are conserved sites bound by antibodies that recognize diverse viral strains, making them attractive targets for vaccine design efforts. Similarly, for a universal influenza vaccine, it would be advantageous to elicit antibodies against conserved sites, like the stem and the 220-loop, which are targeted by broadly cross-reactive antibodies that protect against diverse influenza A viruses.

Various immunogen design strategies have been used to direct immune responses towards subdominant epitopes. Masking unwanted immunodominant sites with glycans or changing their sequence has shown some success^24^. However, these approaches can also inadvertently create new immunogenic regions, may only partially disrupt target sites, or can cause immune responses to shift unpredictably outside the intended epitope^20^. Another strategy involves using subdomains of viral proteins to better expose desired epitopes and remove immunodominant ones, as was done for “headless” HA vaccine candidates that aim to elicit stem-directed antibodies. For the 220-loop epitope, which is partially buried at the interface of HA head units on the trimer, the HA head monomer could serve as an ideal subdomain to increase its exposure in a near native environment. However, as previous observations and our data demonstrate, the HA head contains many other immunodominant sites that are the focus of elicited antibodies and limit responses against the 220-loop. To address this, here we developed epitope scaffold immunogens that display the 220-loop on non-HA derived proteins. These proteins display the target epitope in the conformation recognized by target antibodies, as evidenced by our crystallography studies, have high affinity for multiple broadly cross-reactive antibodies, and lack immunodominant sites encoded by HA.

First-generation 220-loop epitope scaffolds were engineered using previously described computational methods and optimized by *in vitro* evolution. Epitope scaffolds engineered with these methods have two potential limitations: 1) presentation of the grafted epitope in a conformation that is different from that on the native protein, resulting in elicited antibodies that recognize the epitope on the immunogen but do not cross-react with viral proteins, and 2) the presence of immunogenic sites on the scaffold that distract from the epitope. Both these issues affected the immunogenicity of the HA220-4 immunogen as revealed by detailed analysis of the elicited antibodies using Nojima culture. 220-loop immunogens were further optimized using recently developed ML-based protein modeling and design methods, which were integrated into our immunogen design workflow, with applications beyond the current work. To ensure optimal epitope presentation, AlphaFold2 was used to model the Rosetta generated epitope scaffolds and revealed unfavorable epitope conformations that were different than those recognized by target antibodies on the HA head. AlphaFold2 models identified scaffold sites surrounding the epitope that control its orientation. These sites were subsequently optimized by high-throughput display on the surface of yeast, in a process that iterated over four rounds between modeling and experimental optimization. During library screening, it was critical to ensure that the selected epitope scaffolds bind to different 220-loop antibodies to confirm that the grafted epitope, just like HA head, can engage a broad antibody response. The deep scanning mutagenesis of HA220-4 binding to different 220-loop directed antibodies highlighted that these antibodies have different epitope scaffold sequence requirements for binding, even though they recognize and induce the same epitope structure. While D2 H1-1/H3-1 mAb is permissive to HA220-4 sequence changes outside the epitope and residues immediately adjacent, FluA-20 mAb shows a much more restrictive substitution profile, where most of the positions tested had some unfavorable substitutions. Therefore, while AlphaFold2 modeling revealed the lowest energy epitope conformation and suggested ways to alter it, screening with different antibodies can ensure that the desired epitope conformation is readily accessible by target antibodies for high-affinity binding. This is important for vaccine design because we and others have previously shown that binding of multiple target antibodies to engineering immunogens displaying a given epitope is essential for eliciting a polyclonal antibody response against that epitope *in vivo*^35^.

Off-target scaffold responses can distract from target epitopes, as is also observed with native viral proteins. Potential immunogenic sites on HA220-4 were identified by isolating a large number of antibodies elicited by this immunogen using Nojima cultures, followed by modeling of the immunogen-antibody complexes with AlphaFold3. While modeling antibody-antigen interactions remains challenging with AlphaFold3 and other ML-based methods, our goal here was to identify immunogenic scaffold regions at low resolution, and we were less concerned with the atomic details of the molecular interactions. In addition, modeling a large number of antibodies binding to the scaffold ensured that immunogenic sites would be ultimately revealed from multiple simulations, even if some individual models may be inaccurate. This approach is general and can be applied to identify immunogenic regions in a target vaccine candidate by integrating a large number of antibodies isolated in a non-biased fashion, like with Nojima cultures, with antibody-antigen modeling. With expected improvements in modeling antibody structure and interactions in the future, this approach is likely to identify finer structural and sequence details that correlate with immunogenicity. Immunogenic regions on HA220-4 were modified using ProteinMPNN and confirmed by loss of binding of the resulting immunogens, HA220-M4 and HA220-M4T, to antibodies elicited by HA220-4 against the scaffold. This scaffold redesign process does not explicitly consider the immunogenicity of the new surface amino acids; therefore, the ProteinMPNN changes may inadvertently create novel immunogenic sites. Indeed, many isolated antibodies elicited by HA220-M4T still target the scaffold region. Future versions of the epitope grafting protocol will explicitly consider ways to minimize the scaffold immunogenicity. For example, ProteinMPNN sequences can be further down selected to minimize the number of predicted B cell epitopes they contain, or the antibody repertoire elicited by different ProteinMPNN versions of an epitope scaffold can be used to train models that predict the immunogenicity of sequence motifs on a given backbone structure.

Unlike the first generation HA220-4 immunogen, the optimized epitope conformation and scaffold sequence of HA220-M4T elicited epitope-specific antibodies that cross-reacted with HA head or trimer, the major goal of this study. Nevertheless, the antibodies isolated after one immunization with HA220-M4T lacked significant breadth outside the A/H3N2/Hong Kong/1968 strain, the origin of the displayed epitope. This is likely because they had not yet acquired all the somatic mutations necessary for breadth, or that their binding relies on contacts mediated by residues in the epitope that are specific to different influenza strains. Additional maturation of these antibodies may be achieved through additional rounds of vaccination with either HA220-M4T , or, more effectively, with heterologous immunogens that present the epitope in a different molecular context, such as HA or structurally unrelated epitope scaffolds that display a version of the epitope found on other influenza strains, like H1N1 or H5N1. Structural analysis of the HA220- M4T elicited antibodies will reveal their molecular features and may inform vaccination regimens to further mature them towards FluA-20-like breadth. Such regimens will be of interest for the development of a universal influenza vaccine that protects against current and emergent viruses. Taken together, our study illustrates how novel ML-based protein structure prediction and design methods can be applied to control the immunogenicity of engineered proteins and to better focus immune responses on conserved, subdominant epitopes. These general approaches will reveal sequence-structure relationships that control immunodominance hierarchies more broadly and could contribute to the development of vaccines against antigenically variable pathogens like influenza, HIV, or HPV.

## Supplementary Figure Legends

**Supplemental Figure 1.**
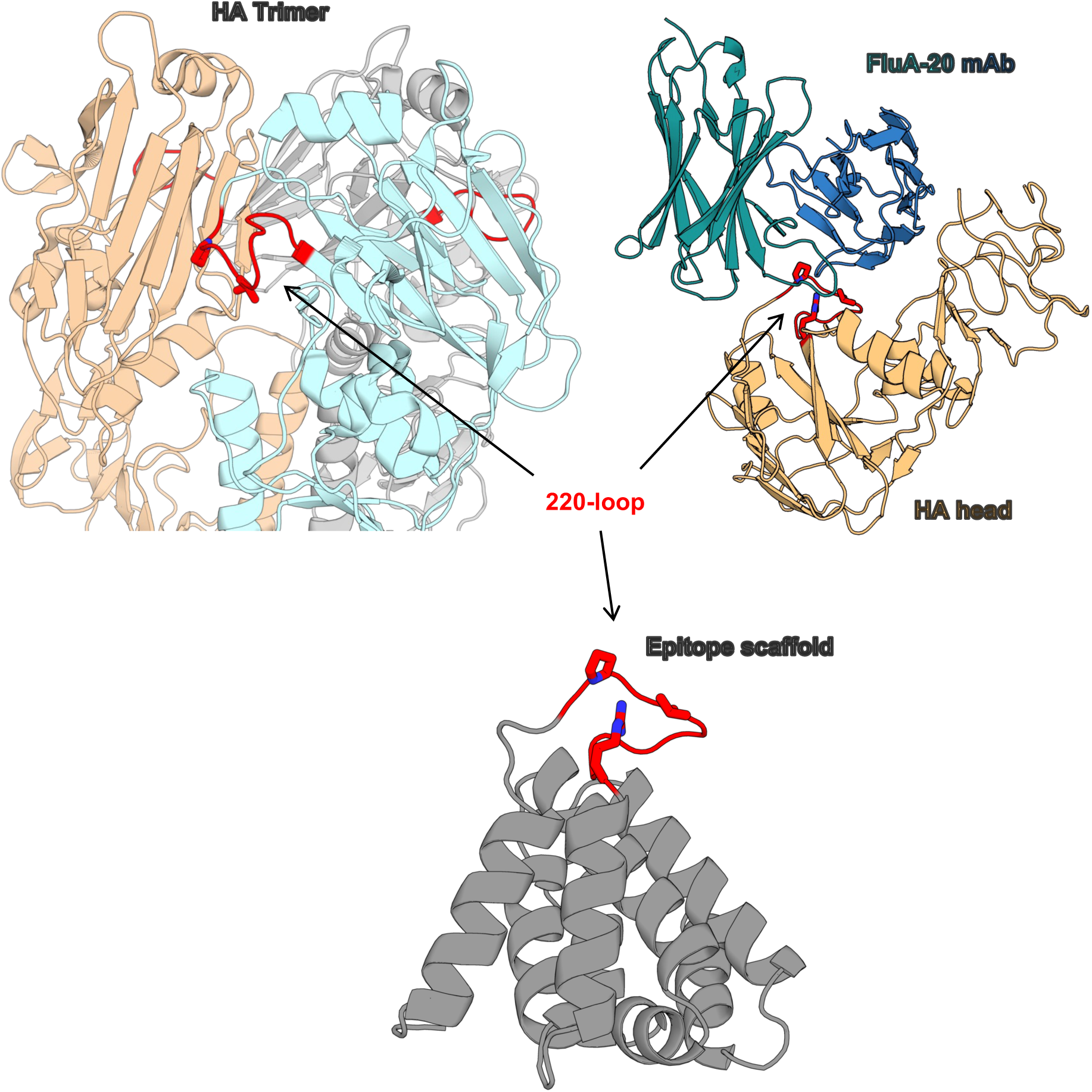
Subdominant, conserved HA epitope targeted by broadly cross-reactive mAbs. (Left) Structure of the HA1 domain of the pre-fusion HA trimer (individual monomers: *grey, pale orange, and pale blue*; A/Hong Kong/1/1968, PDBid:4FNK^38^), with the 220-loop (*red*) highlighted and key epitope residues P221 and V223 (sticks), which are partially occluded on pre-fusion HA. (Right) Structure of HA head (*pale orange,* A/Hong Kong/1/1968) bound to FluA-20 mAb (*green/blue*) (PDBid:6OCB^7^) with key 220-loop epitope residues P221, V223, and R229 (sticks). (Lower) Computational model of HA220 (*grey*) with the grafted epitope shown in *red,* with key 220-loop epitope residues P221, V223, and R229 (sticks) completely exposed.

**Supplemental Figure 2.**
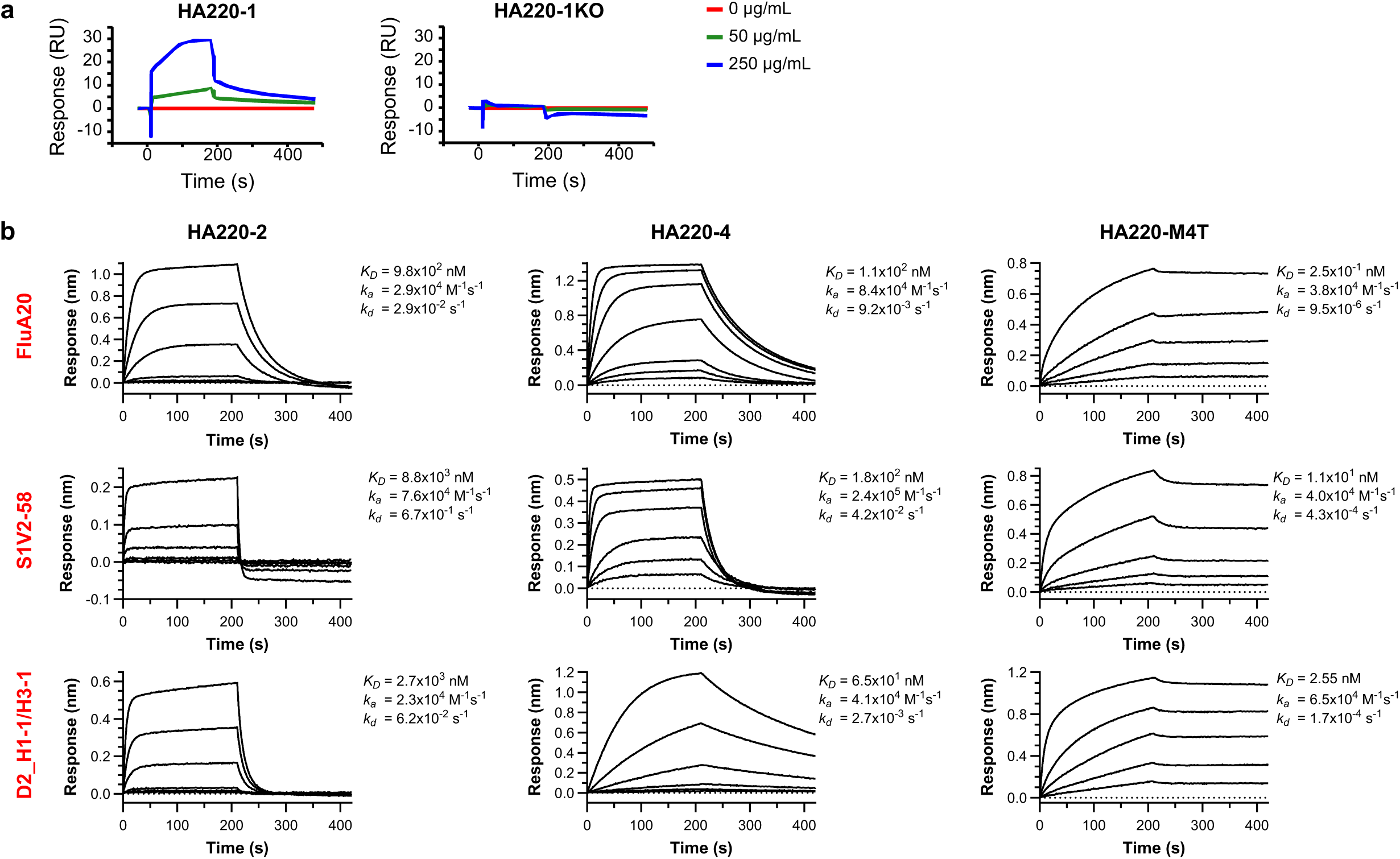
BLI binding curves of HA220 immunogens interacting with target mAbs.

**Supplemental Figure 3.**
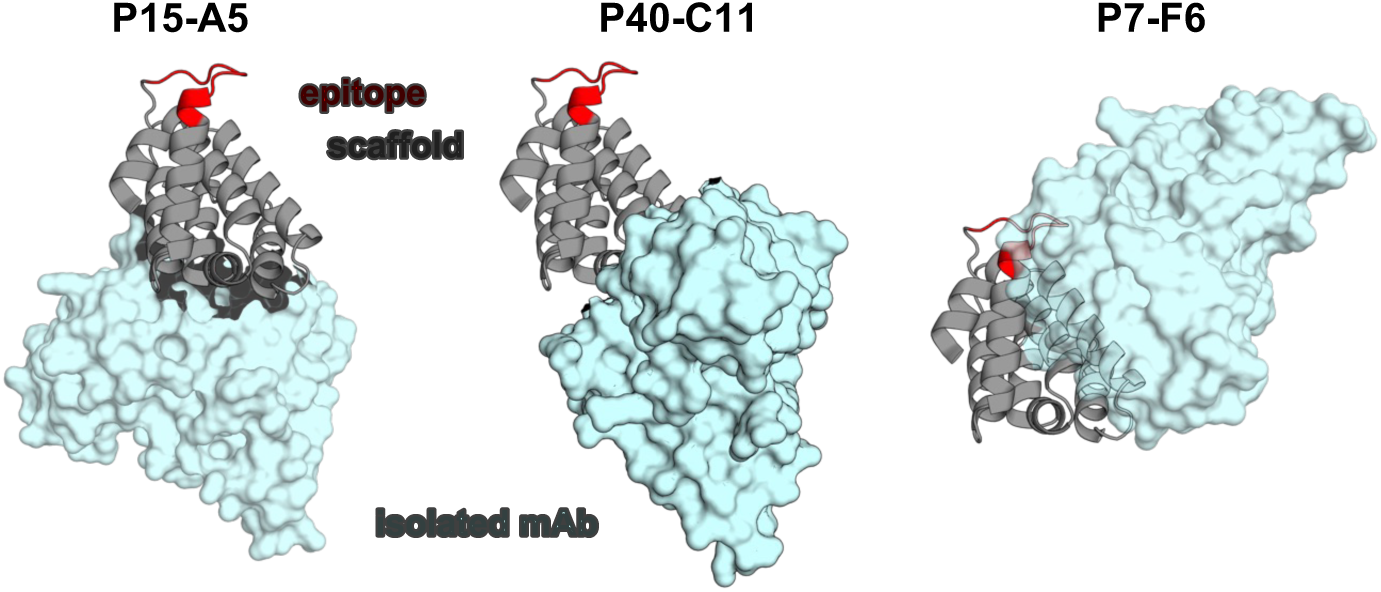
AlphaFold3 models of representative mAbs from *(g)* in complex with HA220-4 (scaffold backbone: *grey*; epitope: *red*; mAb *cyan*)..

**Supplemental Figure 4.**
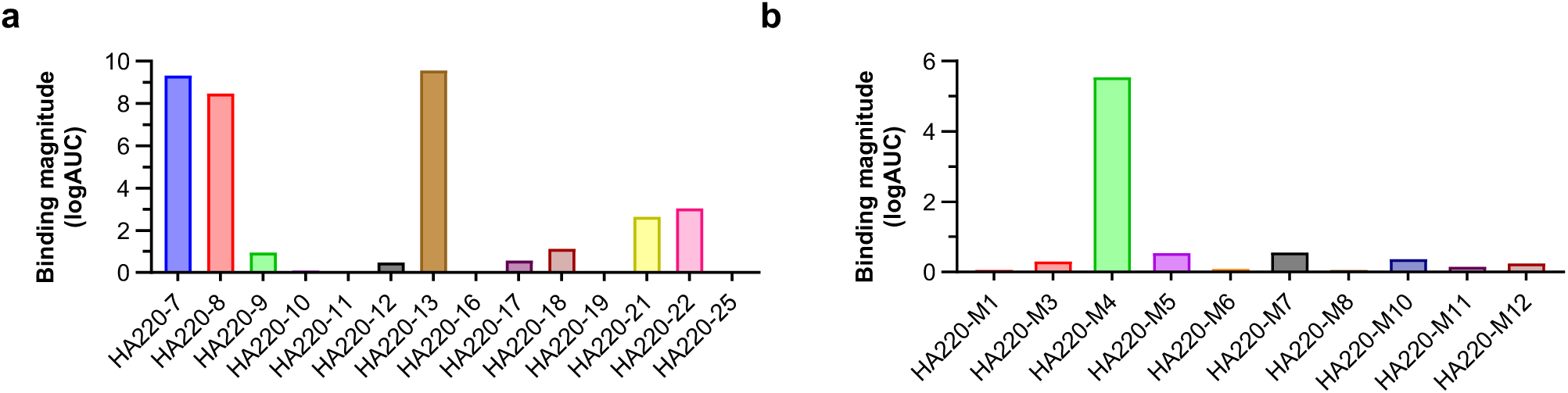
Binding of HA220 scaffolds to mabs measured by ELISA. ELISA binding data of HA220 RS and MPNN immunogens to FluA-20.

**Supplemental Figure 5.**
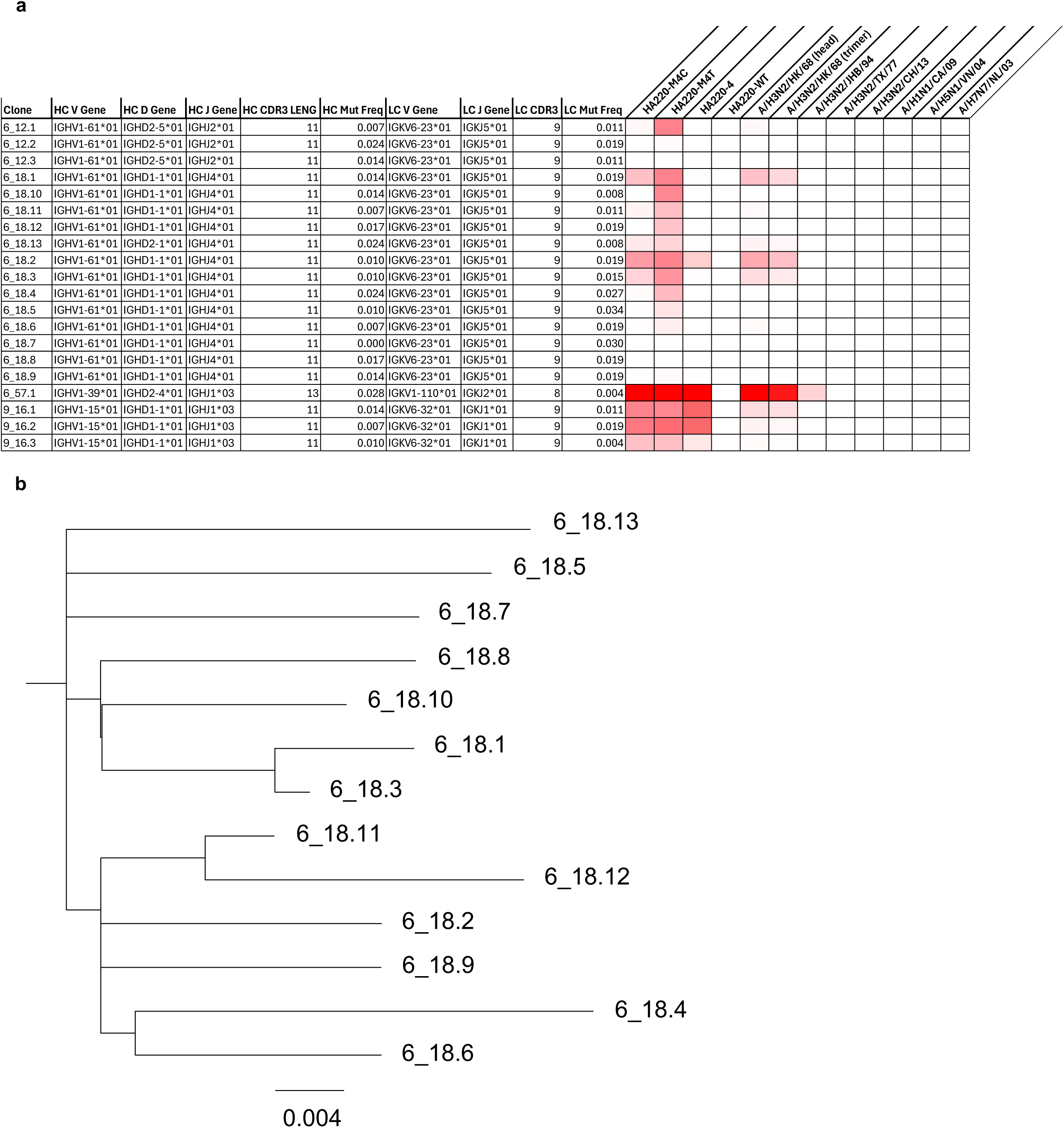
(**A**) Table showing details of HA reactive clones. (B) Tree of 6_18 clone.

**Supplemental Table 1.**

|  | HA220-7 + D2 H1-1/H3-1 H3 Fab | HA220-M4C + S1V2-58 Fab |
| --- | --- | --- |
| *Data collection* |  |  |
| Space group | $P4_122$ | $C2_1$ |
| Cell dimensions |  |  |
| a, b, c (Å) | 110.9, 110.9, 164.8 | 238.6, 158.9, 136.7 |
| $\alpha$, $\beta$, $\gamma$ (°) | 90.0, 90.0, 90.0 | 90.0, 124.9, 90.0 |
| Resolution (Å) | 82.43-2.40 (2.49-2.40) | 71.11-3.90 (4.07-3.90) |
| $R_{\text{merge}}$ | 0.188 (1.931) | 0.243 (0.957) |
| $I/\sigma I$ | 9.7 (1.6) | 4.2 (1.7) |
| CC1/2 | 0.997 (0.780) | 0.931 (0.357) |
| Completeness (%) | 100.0 (100.0) | 99.7 (99.9) |
| Redundancy | 12.8 (12.4) | 3.3 (3.5) |
| *Refinement* |  |  |
| $R_{\text{work}}/R_{\text{free}}$ (%) | 21.5/24.3 | 21.7/27.6 |
| No. atoms |  |  |
| Protein | 4466 | 18082 |
| Ligand | 0 | 0 |
| Water | 130 | 0 |
| Average B-factors |  |  |
| Protein | 61.3 | 112.0 |
| Solvent | 55.6 |  |
| R.m.s deviations |  |  |
| Bond lengths (Å) | 0.003 | 0.003 |
| Bond angles (Å) | 0.58 | 0.69 |
| Clashscore | 2.59 | 4.76 |
| Ramachandran |  |  |
| Favored (%) | 97.7 | 95.1 |
| Allowed (%) | 2.3 | 4.6 |
| Outliers (%) | 0.0 | 0.3 |

## Materials and Methods

### Computational design of epitope scaffolds

#### Design of HA220 epitope scaffolds by backbone grafting

A database of 9884 candidate scaffolds^35^ was created by selecting from the Protein Data Bank (PDB) structures that were: 1) determined by x-ray crystallography; 2) high resolution (<2.8 Å); 3) monomeric; 4) expressed in *E. coli*; 5) that did not contain ligands. Epitope scaffolds that display the stem helix epitope recognized by the FluA-20 antibody (PDBid: 6ocb) were designed by backbone grafting as previously reported and using the RosettaScripts described by Silva et. al. The same set of curated parent scaffolds as the one used for side chain grafting above was searched here. The structure of the 220-loop fragment ^220^RPWVRGLSSR^229^ from PDBid:7sjs was grafted. The identity of residues V224 through R229 were kept fixed during the design process while the other epitope residues were allowed to change. The following parameters were used:

RMSD_tolerance="2.0" NC_points_RMSD_tolerance="2.0"

clash_score_cutoff="5" clash_test_residue="GLY"

hotspots="4:10"

combinatory_fragment_size_delta="3:0”

max_fragment_replacement_size_delta="-8:8"

full_motif_bb_alignment="0"

allow_independent_alignment_per_fragment="0"

graft_only_hotspots_by_replacement="0"

only_allow_if_N_point_match_aa_identity="0"

only_allow_if_C_point_match_aa_identity="0"

revert_graft_to_native_sequence="0"

allow_repeat_same_graft_output="0" />

The initial hits were visually examined to ensure appropriate epitope transplantation and antibody-scaffold interaction. Additional changes were introduced in the candidate epitope scaffold using Rosetta fixed backbone design to remove antibody-epitope scaffold contacts that were not due to the target epitope and to ensure proper interactions between the epitope and the rest of the scaffold.

#### ProteinMPNN design

For ProteinMPNN redesign, the Rosetta model was entered into ProteinMPNN at the website, https://huggingface.co/spaces/simonduerr/ProteinMPNN using a sampling temperature of 0.1 and backbone noise of 0.02 and the either whole epitope sequence and N69 was fixed, or just the P221, V223 and R229 (HA numbering) were fixed. ES structures were modelled using the integrated Colabfold prediction and scored by RMSD and pLDDT, and the highest scoring were re-predicted using Alphafold2.

#### *In-silico* prediction of beneficial 2-, 3-, 4- and 5- point mutations

We sought to identify 20 beneficial higher-order mutations (5 sequences each for 2-, 3-, 4- and 5-point mutations) using our single-point mutation data from HA220-4. This posed a significant computational challenge due to epistatic effects, where mutations are non-additive—two individually beneficial mutations may not produce an additive benefit when combined. To account for epistasis, we used Protein Language Models (PLMs) to identify multi-point mutation targets.

PLMs, trained on millions of protein sequences, capture the statistical patterns underlying protein structure and function, enabling them to predict epistatic effects with considerable success^39^. For robust selection, we employed three different PLMs in a consensus approach: ESM-2 650M as a general-purpose PLM^40^, ESM-1v for its training on the broader UniRef90 database, and Raygun (doi:10.1101/2024.08.13.607858) because its fixed-length representations have been shown to effectively capture protein structural folds. Each PLM was separately trained on our HA220-4 single-mutation data, enabling inference of antibody affinities from sequence alone.

For each mutation category (N=2, 3,4, 5) we produced 1,500 candidates, totaling 6,000 candidate sequences across all four categories. For 2-point mutations, we exhaustively enumerated all feasible combinations, selecting the top 1,500 candidates ranked by their single-point phenotype scores. However, exhaustive enumeration becomes computationally intractable for higher-order mutations and produces candidates with limited sequence diversity. For 3-, 4-, and 5-point mutations, we therefore generated candidates through importance sampling: mutations were probabilistically selected based on their single-point phenotype scores, with the constraint that selected mutations could not occur at the same residue position.

For each mutation order (N=2,3,4,5), we filtered candidates using all three PLMs, retaining only those ranked within the top 200 by all three models. From these high-confidence candidates, we selected the 5 most sequence-diverse variants by K-Medoid clustering using distances derived from Raygun’s fixed-length representations, yielding 20 total candidates for experimental validation.

### Plasmids and DNA synthesis

Genes encoding designed epitope scaffolds were commercially synthesized and cloned into pET29b (Genscript and Twist Bioscience). Genes encoding the antibody heavy and light chains were similarly synthesized and cloned into the pcDNA3.1 vector (GenScript). Oligonucleotides were synthesized by IDT. Mutations were introduced in the synthesized plasmids by Q5 mutagenesis (NEB).

### Recombinant protein expression and purification

#### Epitope scaffolds

Plasmids encoding epitope scaffolds were transformed into *E. coli* BL-21 (DE3) (New England Biolabs) cells and 5 mL starter cultures were grown overnight at 37□°C in Lysogeny Broth (LB) supplemented with 50 μg/mL kanamycin. The cultures were diluted 1:100 in Terrific BrothTB media (RPI) supplemented with kanamycin and grown at 37□°C to an OD600 of ∼0.6. The temperature was subsequently lowered to 16□°C and the cultures were grown to OD600 of ∼0.8 and induced with isopropyl β-D-1-thiogalactopyranoside (IPTG) to a final concentration of 0.5 mM. The cultures were shaken for 16-18 hours at 200 rpm. Cell pellets were collected by centrifugation at 14,500xg and lysed in B-PER Reagent (ThermoFisher Scientific) according to the manufacturer’s protocol. The lysates were centrifuged at 14,000xg for 30 minutes at 4□°C. The supernatant was incubated with Ni-NTA beads (Qiagen) equilibrated with native binding buffer (20 mM NaH2PO4, 500 M NaCl, 2% glycerol, 10 mM imidazole, pH=7.5) for an hour at 4□°C. The beads were settled by centrifugation, and the supernatant was removed by pipetting. The beads were washed with wash buffer (50 mM NaH2PO4, 500 M NaCl, 2% glycerol, 30 mM imidazole, pH=7.5) after which the protein was eluted with elution buffer (20 mM NaH2PO4, 500 M NaCl, 250 mM imidazole, pH=7.5). Protein expression and purity was confirmed by SDS-PAGE analysis and quantified spectrophotochemically at 280 nm on a Nanodrop 2000 (ThermoFisher Scientific). The eluted protein was concentrated on 3 kDa MW spin columns (ThermoFisher Scientific) and further purified by size exclusion chromatography in 20 mM NaH2PO4, 150 mM NaCl, pH=7.5, on an AKTA-Go FPLC (Cytiva) using a Superdex 200 Increase 10/300GL column. Fractions containing monomeric protein were pooled and concentrated as above.

#### Monoclonal antibodies

Antibodies were produced as previously described^41^. Expi293f cells (Invitrogen) were split on the day of transfections at a density of 2.5×10^6^ cells/mL with an average of 94-99% viability. The cells were then transiently transfected with an equimolar amount of heavy and light chain plasmids using expifectamine (Invitrogen) incubated in Opti-MEM (Invitrogen). The morning post transfection, enhancers were added to the cultures, and cultures were harvested after five days. The cells were spun down by centrifugation, and the supernatant was filtered using a 0.8 μm filter. The supernatant was then added to equilibrated Protein A beads and incubated for an hour at 4 °C. The beads were then separated from the supernatant and washed with a buffer containing 20 mM Tris and 350 mM NaCl at pH=7 in a column. The antibodies were then eluted from the beads with a 2.5% glacial acetic acid elution buffer, and was buffer exchanged into 25 mM Citric Acid, 125 mM NaCl buffer at pH=6. Antibodies were then further quality checked for purity and expression using SDS-PAGE analysis and quantified by measuring absorbance at 280 nm (Nanodrop 2000).

### Optimization of HA220 epitope scaffolds by yeast display

#### Library design and synthesis

Combinatorial mutagenesis libraries were synthesized on a BioXp system (CodexDNA) or were synthesized by Twist Bioscience. SSVL libraries were synthesized by Twist Bioscience.

Libraries were then amplified with Q5 polymerase and purified by agarose gel extraction and PCR cleanup (Qiagen) as per the manufacturer’s protocol.

#### Library transformation into S. cerevisiae

*S. cerevisiae* EBY100 yeast cells were transformed to express HA220 libraries as previously described^42–44^. Cells were transformed by electroporation with a 3:1 ratio of 12 µg HA220 library DNA and 4 µg pCTCON2 plasmid digested with BamHI, SalI, NheI (NEB). The typical sizes of the transformed libraries, determined by serial dilution on selective plates, ranged from 2-6×10^7^. Between 60% and 80% of the sequences recovered from the transformed libraries were confirmed to contain full length, in-frame genes by Sanger sequencing (Genewiz). Yeast libraries were grown in SDCAA media (Teknova) supplemented with pen-strep at 30 °C and shaking (225 rpm).

#### Library screening by FACS

HA220 expression on the surface of yeast was induced by culturing the libraries in SGCAA (Teknova) media at a density of 1×10^7^ cells/mL for 24-36 hours. Cells were washed twice in ice cold PBSA (0.01 M sodium phosphate, pH=7.4, 0.137M sodium chloride, 1 g/L bovine serum albumin) and incubated for 1 hour at 4□°C with varying concentrations of 220-loop mabs. Cells were then washed twice with PBSA, resuspended in secondary labeling reagent α c-myc:FITC (ICL) and anti-human:R-PE (Sigma-Aldrich) and incubated at 4 °C for 30 minutes. Cells were washed twice with PBSA after incubation with the fluorescently labeled probes and sorted on a Sony Sorter. Double positive cells for PE and FITC were collected and expanded for one week in SDCAA media supplemented with pen-strep before DNA isolation from selected clones. FACS data was analyzed with Flowjo_v10.6 software (Becton, Dickinson & Company). All clones selected by FACS were expanded, and their DNA was extracted (Zymo Research) for analysis by Next Generation Sequencing (Illumina) and Sanger sequencing (Genewiz).

#### Sequence analysis of HA220 libraries

HA220 encoding plasmids were recovered from yeast cultures by miniprep with the Zymoprep yeast plasmid miniprep II kit (Zymo Research) as previously described^44^. Isolated DNA was transformed into NEB5α *E. coli* (NEB), and the DNA of individual bacterial colonies was isolated (Wizard Plus SV Minipreps, Promega) and analyzed by Sanger sequencing. To prepare for Next Generation Sequencing, the HA220 SSVL insert from isolated plasmids was amplified by PCR using Q5 polymerase (NEB). DNA samples were prepped and run using the Illumina MiSeq v3 reagent kit following manufacturer’s protocols. Illumina sequencing returned an average of 21.6 million reads per sample, of which an average of 20.7 million mapped to the scFv amplicon. Sequencing data was processed using Geneious Prime and in-house scripts to compute the amino acid frequency and distribution.

### Structural analysis by x-ray crystallography

D2 H1-1/H3-1 Fab was mixed with a 1.5-fold molar excess of RS7 scaffold and S1V2-58 Fab was mixed with a 1.5-fold molar excess of HA220-M4C scaffold to bind overnight at 4 °C. Intact complexes were purified using a HiLoad 16/600 Superdex200 size-exclusion column (Cytiva) in 2 mM TRIS pH=8.0, 200 mM NaCl, 0.02% NaN_3_. SEC-purified complexes were concentrated to 7.00 mg/mL and used to prepare high-throughput sitting drop crystallization screens using an Oryx4 liquid handler (Douglas Instruments). HA220-7 + D2 H1-1/H3-1 crystals grew in a mother liquor composed of 0.1 M HEPES pH=7.5 and 25% PEG 3350. Crystals of HA220-M4C + S1V2-58 grew in a mother liquor composed of 0.1 M BIS-TRIS pH=5.5 and 2.0 M ammonium sulfate. Single crystals of each complex were cryoprotected with mother liquor supplemented with 20% glycerol prior to manual plunge freezing in liquid nitrogen.

Diffraction data for both complexes were collected at NSLS2 Beamline 19-ID. Data were indexed and integrated in iMOSFLM^45^ prior to merging and scaling in AIMLESS^46^. Phases were solved by molecular replacement using PhaserMR^47^ with PDB IDs: 6XPR chains B and C (D2 H1-1/H3-1) and 6XPY chains B and C (S1V2-58) as search ensembles. AlphaFold3^32^ was used to generate molecular replacement ensembles for the HA220-7 and HA220-M4C scaffolds. Coordinates were then iteratively refined using Coot^48^, Phenix^49^ and ISOLDE^50^. Crystallographic data collection and refinement statistics are summarized in **Supplementary Table 1**. Images of structures were produced using Pymol (Schrodinger) and RMSDs were calculated using ChimeraX^51^.

### Antigenic characterization of epitope-scaffolds

#### Antibody binding by ELISA

Enzyme-linked immunosorbent assays (ELISAs) were performed as described previously^41^. The following proteins were used, either produced as above or available commercially: HK/68 trimer (Sino Biologicals, #40715-V08H); HK/68 head; HA220-1; HA220-2; HA220-4; HA220-4KO; HA220-7; HA220-7KO; HA220-4-SV1; HA220-4-SV2; HA220-4-SV3; HA220-M4; HA220-M4C; HA220-M4T; HA220-M4T^+glyc^; A/H3N2/TX/77; A/H3N2/BK/79; A/H3N2/JB/94; A/H3N2/TX/12; A/H3N2/CH/13; A/H1N1/CA/09; A/H5N1/VN/04; A/H5N1/IM/05; A/H7N7/NL/03. HA proteins were coated overnight on 384-well plates (Corning, #3700) at 4 °C. Purified antibodies were used at 100 μg/mL starting concentrations and were then serially diluted 3-fold and incubated at room temperature for 1 hour. Goat anti-human IgG-HRP secondary antibody (Jackson ImmunoResearch Laboratories, #109-035-098) was diluted 1:15,000 and added to plates, incubated for one hour, and developed using TMB substrate (SureBlue Reserve, KPL, #5120-0083). Plates were read at 450 nm on a Cytation 1 plate reader (BioTek). Data was analyzed and plotted using GraphPad Prism version 10.6.1 for Windows (GraphPad Software, Boston, Massachusetts USA, www.graphpad.com).

#### Binding by Surface Plasmon Resonance (SPR) / Biolayer Interferometry (BLI)

A Sartorius Octet R4 instrument was used to measure binding of HA220 variants to a panel of antibodies (FluA-20, S1V2-58, D2 H1-1/H3-1). All the assays were performed in 1X HBS-EP Buffer (Cytiva) supplemented with 0.1% BSA. The final volume for all the solutions was 200 μl per well. Assays were performed at 25 °C in solid black tilted-bottom 96-well plates (Griener). Human antibodies (2.5 μg/ml) in 1X HBS-EP buffer were used to load Octet ProA Biosensors (Sartorius) for 90 seconds. Biosensor tips were then equilibrated for 60 seconds in 1X HBS-EP, 0.1% BSA buffer before binding assessment. Binding of HA220 variants were allowed for 210 seconds and dissociation was performed for 420 seconds. Data analyses were carried out using Octet Analysis Studio software, version 13.0.

### Mouse immunization studies to quantify sera responses

#### Immunization studies in BALB/c mice

C57BL/6 mice were obtained from the Jackson Laboratory. Eight to 12-week-old female mice were used in this study. All mice were maintained under specific pathogen-free conditions at the Duke University Animal Care Facility. All experiments involving animals were approved by the Duke University Institutional Animal Care and Use Committee.

For some experiments, mice were immunized with 20 μg of immunogens (see below) in the footpad of the right hind leg. Two weeks (14-15 days) after immunization, GC B cells were sorted from popliteal LN for single-cell Nojima cultures (see below). All antigens were mixed with Alhydrogel^®^ adjuvant 2% (final 1%) before immunizations.

For some experiments, mice were primed intraperitoneally with 20 μg of HA H3 Hong-Kong 1968 head monomer immunogen and boosted twice with Alhydrogel^®^ adjuvant (alum-only) or with HA220 immunogen intraperitoneally. Immunizations were performed 4 weeks apart. Serum samples were collected 1 week after each immunization. All antigens were mixed with Alhydrogel^®^ adjuvant 2% (final 1%) before immunizations.

#### Sera binding by ELISA

ELISA binding assays with mouse sera were performed as described previously^41^. 384-well plates were prepared in the same manner as in *Antibody binding by ELISA*. Mouse sera were diluted 1:100 while antibody controls were diluted to 100 μg/mL and serially diluted 5-fold. Samples were added and incubated at room temperature for 1.5 hours. The corresponding IgG-HRP secondary antibody was diluted and subsequently added (goat anti-human IgG-HRP, 1:15,000 dilution, Jackson ImmunoResearch Laboratories, #109-035-098; or goat anti-mouse IgG-HRP, 1:8,000 dilution, SouthernBiotech, #1030-05). After 1 hour incubation, plates were developed using TMB substrate (SureBlue Reserve, KPL, #5120-0083) and read at 450 nm on a SpectraMax Plus plate reader (Molecular Devices). Data was analyzed and plotted using GraphPad Prism version 10.6.1 for Windows (GraphPad Software, Boston, Massachusetts USA, www.graphpad.com).

### Mouse immunization studies for Nojima analysis

#### Flow Cytometry

Germinal center (GC) B cells (GL-7^+^B220^hi^CD38^lo^IgD^-^CD138^-^) in popliteal LNs of immunized mice were identified as described^52–54^. Labeled cells were analyzed/sorted in a FACSymphony^TM^ S6 Cell Sorter with DIVA option (BD Bioscience). Flow cytometric data were analyzed with FlowJo software (Treestar Inc.). Cell doublets and dead cells (propidium iodide incorporated cells) were excluded from our analyses^52–54^. mAbs used in this study: FITC-conjugated anti-GL-7 (GL7, BD Bioscience), BV786-conjugated anti-B220 (RA3-6B2, Biolegend), PE-Cy7-conjugated anti-CD38 (90, BioLegend), BV510-conjugated anti-IgD (11-26c.2a, BioLegend), BV605-conjugated anti-CD138 (281-2, BD Bioscience).

#### Single-cell Nojima culture

GC B cells were expanded in single cell cultures^52–54^. Briefly, single B cells were directly sorted into each well of 96-well plates that contained NB-21.2D9 feeder cells (2,000 cells/well) and recombinant mouse IL-4 (Peprotech; 2 ng/ml) in B cell media (BCM); RPMI-1640 (Invitrogen) supplemented with 10% HyClone FBS (Thermo scientific), 5.5×10^-5^ M 2-mercaptoethanol, 10 mM HEPES, 1 mM sodium pyruvate, 100 units/ml penicillin, 100 μg/ml streptomycin, and MEM nonessential amino acid (all Invitrogen). Two days after sorting, 50% (vol.) of culture media were removed from cultures and 100% (vol.) of fresh BCM were added to the cultures. Two-thirds of the culture media were replaced with fresh BCM every day from day 4 to day 8. On day 10, culture supernatants were harvested for ELISA determinations and culture plates that contained clonally expanded single B cells were stored at −80 °C for V(D)J amplifications.

#### ELISA and Luminex assays

Presence of total and antigen-specific IgG in serum samples and in culture supernatants were determined by ELISA and Luminex multiplex assay^52–54^. Diluted culture supernatants (1:10 in PBS containing 0.5% BSA and 0.1% Tween-20) were screened for the presence of IgGs by standard ELISA. IgG^+^ samples were then screened for binding to immunogen antigens and influenza hemagglutinins by Luminex assay. Briefly, serum samples or culture supernatants were diluted (initial dilutions of 1:100, and then 3-fold, 11 serial dilutions for serum samples; 1:10 or 1:100 for culture supernatants) in 1xPBS containing 1% BSA, 0.05% NaN3 and 0.05% Tween20 (assay buffer) with 1% milk and incubated for 2 hours with the mixture of antigen-coupled microsphere beads in 96-well filter bottom plates (Millipore). After washing with assay buffer, these beads were incubated for 1 hour with PE-conjugated goat anti-mouse IgG Abs (Southern Biotech). After three washes, the beads were re-suspended in assay buffer, and the plates were read on a Bio-Plex 3D Suspension Array System (Bio-Rad). The following antigens were coupled with carboxylated beads (Luminex Corp): BSA (Affymetrix), goat anti-mouse Igκ, goat anti-mouse Igλ, goat anti-mouse IgG (all Southern Biotech), a panel of HAs and engineered antigens, including HK/68 head, A/H1N1/CA/09, A/H3N2/CH/13, HA220-KO, HA220-4, HA220-M4T^glyc^ (all in house), and HK/68 trimer (SinoBiological).

#### Serum competition assay

S5V2-29 inhibition assay was carried out as described^8^ with modification. Briefly, diluted (1:100) mouse sera were incubated overnight at 4 °C with Luminex beads coupled with HA H3 HK-68 or with anti-human IgG. Beads were then incubated for 2 hours at room temperature with 10 ng/ml of recombinant human S5V2-29 Ab^8^. After washing plates, beads were incubated for 1 hour with PE-conjugated mouse anti-human IgG Abs (JDC-10; Southern Biotech). We calculated relative MFI values (MFI on HA H3 HK-68/ MFI on anti-human IgG) for samples. Average of the relative MFI values from pre-immune sera (n=48) that did not contain HA H3 HK-68 binding IgGs were set as 100% (no inhibition) for screening samples against HA H3 HK-68.

#### Immunogenetic annotation of the PACBIO sequencing data

Consensus heavy and light sequences for each well were processed using an updated version of the Absolute program. In some cases, more than one sequence for the same chain was returned for a well. In these cases, only the sequences with more than 10 support reads were considered for the next processing steps. If all sequences have the same V allele, then the sequence with the most consensus reads is kept. If multiple V genes/alleles are present, then all functional (modulo 3 CDR3, conserved cysteines and tryptophan/phenylalanine, and modulo 3 insertions and deletions) and productive (no stop codon) sequences are kept. Immunogenetic annotation of the sequences (V, D, and J segment assignment, CDR3 length, V mutation frequency) was performed using the Partis program^55^ with the Partis mouse reference genes from the C57BL6 mouse strain for the VH segment, and the default partis mouse library for the D, VK, VL. and and all J segments. Clonal partitioning was done on the heavy chain sequences using the Cloanalyst program^56^. Wells that had one in frame heavy chain sequence and at least one kappa or lambda in frame rearrangement were considered in the subsequent analysis. For the sequences with more than one light rearrangement, if the well by heavy chain was part of an expanded clone, the light chain that matched the majority light chain of that clone was chosen. Paired chain clonal tree inference including UCA inference was done with the Cloanalyst program [56].

## Acknowledgments

We would like to acknowledge the following people for their efforts and contributions to this work: James San, Joshua Martin Beem, Gracie Li, and Irina Dumbravanu (Duke University), Chiaki Nishihara (Tokyo University of Agriculture and Technology), Ryo Saotome (North Carolina School of Science and Math), Shrina Shah (North Carolina School of Science and Math). K.D. and R.S. acknowledge the support of the Whitehead Scholarship at Duke University. The NIH funded this research through grants R01AI155804 to M.L.A.; and 1U54AI191253 and 1U01AI186999 to R.S.

## Author contribution

Experimental work was carried out by D.J.M., A.W., A.B.K., C.H., D.W., J.C., P.L., B.P., B.R., M.F., K.W., H.B., P.K., X.L., and M.K. Crystallographic studies - D.W. Nojima immunization and B-cell isolation studies - A.W. P.L. B.P. X.L. G.K. M.K. Computational immunogen design and optimization - D.J.M., K.D., R.S., and M.L.A. Sequence analysis of isolated antibodies – E.V.L. and K.W. The study was directed by M.L.A, M.K., and G.K. The manuscript was written by D.M., C.H., and M.L.A., with input from M.K., R.S., D.W., and G.K.

